# MOSAIC: A scalable framework for fMRI dataset aggregation and modeling of human vision

**DOI:** 10.64898/2025.11.28.690060

**Authors:** Benjamin Lahner, Mayukh Deb, N. Apurva Ratan Murty, Aude Oliva

**Affiliations:** Computer Science and Artificial Intelligence Laboratory, MIT, Cambridge, MA, USA; Department of Ophthalmology, Byers Eye Institute, Stanford University School of Medicine, Stanford University, Stanford, CA, USA; Stanford Bio-X, Stanford University, Stanford, CA, USA; Wu Tsai Neurosciences Institute, Stanford University, Stanford, CA, USA; Cognition and Brain Science, School of Psychology, Georgia Tech, Atlanta, GA, USA; Computational Cognition, Georgia Tech, Atlanta, GA, USA

**Keywords:** fMRI, Encoding Models, Decoding Models, functional localizer

## Abstract

Recent large-scale vision fMRI datasets have been invaluable resources to the vision neuroscience community for their deep sampling of individual subjects and diverse stimulus sets. However, practical limitations to the number of subjects, stimuli, and trials that can be collected prevent individual fMRI datasets from reaching the scale necessary for modern modeling approaches and robust conclusions. Here, we introduce MOSAIC (Meta-Organized Stimuli And fMRI Imaging data for Computational modeling), a fMRI dataset aggregation framework designed to leverage the richness of individual datasets for computationally intensive modeling and robust tests of generalization. MOSAIC is composed of eight large-scale vision fMRI datasets totaling 93 subjects, 430,007 fMRI-stimulus pairs, and 162,839 naturalistic and artificial stimuli. A shared fMRI preprocessing pipeline and a filtered test-train split minimizes dataset-specific confounds and test-set leakage when aggregating the datasets. Crucially, additional datasets can be integrated into MOSAIC post hoc, allowing MOSAIC to evolve according to the community’s interests. We use MOSAIC to show that perceptually diverse stimulus sets consistently improve decoding accuracy and stability, carrying implications for future fMRI stimulus set design. We then jointly train brain-optimized encoding models across subjects and datasets to predict fMRI activity of all visual cortex and even the whole brain. In silico functional localizer experiments performed on these digital twin models can recover subject-specific category-selective cortical regions, thereby validating our approach. Together, MOSAIC provides a scalable and community-driven solution to build robust, larger-scale models of human vision.

## 1 Introduction

Naturalistic experimental paradigms are essential to build models of real-world brain and behavioral responses [1–6]. Large-scale fMRI datasets are central to this effort as their increased number of subjects, number of stimuli, and stimulus diversity make them especially well-suited for data-driven machine learning methods to uncover latent neural structure under more ecological contexts [1, 7–13]. However, the size of individual fMRI datasets is quickly approaching its upper limit while this promising effort of naturalistic experimental research is still in its infancy [11, 14]. Worse still, the space of ecological contexts is too vast for a single research group to cover; for example, even the most massive image-based fMRI dataset cannot capture neural responses to dynamic videos [15–17], which may in turn be of insufficient duration to capture responses uniquely evoked by long form videos [18–21], ad infinitum. The neuroscience community in aggregate, however, has collected vast amounts of fMRI data across a range of ecological contexts, but the field lacks a framework to unify them.

Despite a growing ecosystem of public fMRI datasets [22–24], datasets are often used in isolation from one another. Differences in preprocessing pipelines, experimental protocols, acquisition systems, registration spaces, and subjects contribute sources of variability in the signal that complicate cross-dataset analyses [25–30]. Additionally, because different datasets may use identical or highly similar stimuli, it is essential for modeling efforts to ensure test-train splits are still independent after aggregating multiple datasets. Overcoming these challenges will allow researchers to simultaneously leverage multiple datasets for enhanced statistical power, stimulus diversity, and generalizability – ultimately leading to a more robust understanding of the human brain.

Recent data collection and modeling efforts have recognized the need of vast amounts of fMRI data to perform large-scale model training or leverage held-out test sets [13, 31–36]. However, their approach of pooling each fMRI dataset’s originally released preprocessed data creates a trade-off between scale and scientific validity. By mixing preprocessing pipelines, brain registration templates, and data formats (e.g., time series and beta estimates), such aggregate datasets introduce new sources of variance that can mask or mimic true neural signals [25–30]. While operating in latent space can bypass issues of topographical brain alignment, it sacrifices the fine-grained, voxel-level interpretability necessary for mechanistic insights. Furthermore, naive aggregation risks unintended test set leakage when stimuli overlap between datasets, rendering claims of generalization invalid. As such, the field currently lacks a robust framework that can achieve massive scale datasets without sacrificing methodological rigor.

To overcome these challenges, we introduce MOSAIC (Meta-Organized Stimuli and fMRI Imaging data for Computational modeling), our blueprint for a community-driven fMRI dataset aggregation framework and dataset that supports generalizable research and data-intensive computational modeling in vision neuroscience. This first version of the MOSAIC dataset has eight large-scale vision fMRI datasets totaling 93 subjects, sufficient for high-powered and reproducible analyses [37–39]. MOSAIC data underwent a shared preprocessing pipeline (fMRIPrep [40] and GLMsingle [41]) and has 430,006 high-quality single trial fMRI-stimulus pairs across 162,839 unique stimuli. The stimulus set is subsequently curated into naturalistic (e.g., real-world images and videos) train, naturalistic test, and artificial (e.g., shapes, noise patterns, gratings) test splits for robust tests of generalizability. MOSAIC has been designed to be an extensible, community-driven resource - datasets of any size can be integrated post-hoc, and the common preprocessing steps can evolve with advancements in experimental paradigms and methodologies.

Here we detail MOSAIC’s preprocessing procedure and showcase how it can be used to both replicate results across subjects and pool data to achieve massive scale for computational modeling efforts. We observe that MOSAIC’s preprocessing pipeline preserves meaningful biological signal across different experimental paradigms and scanners. Next, we find that perceptually diverse stimulus sets significantly improve decoding accuracy and stability, further motivating MOSAIC’s goal of compiling a diverse aggregate stimulus set and providing actionable recommendations for future fMRI datasets to best sample their stimulus set. Finally, we jointly train brain-optimized neural networks across the MOSAIC aggregate dataset and find that we are able to directly predict subject-specific brain activity better than corresponding subject-specific models, thus overcoming a major hurdle in scaling fMRI data. We go beyond reporting correlational model encoding accuracy by demonstrating the scientific value of these brain optimized models as in silico data generators that can recover subject-specific category-selective functional regions. In this light, MOSAIC’s flexibility will enable the neuroscience community to continue collecting datasets to target research questions that matter to them most while simultaneously allowing these datasets to be part of an aggregate collection to benefit large-scale modeling efforts and encourage reproducible and generalizable results. With MOSAIC, the computational cognitive visual neuroscience community can join forces to build a collaborative, scalable, and generalizable data foundation for the continual advancement of our collective understanding of human vision.

## 2 Results

### 2.1 MOSAIC aligns fMRI datasets

#### 2.1.1 Constructing MOSAIC

The construction of MOSAIC followed a three-step process. *First*, we systematically evaluated candidate fMRI datasets and selected only those that met 3 key criteria: they 1) used vision-only paradigms, 2) employed event-related fMRI task paradigms, and 3) included perception-related behavioral tasks. This process led us to identify 8 open-access fMRI datasets (Fig. 1). *Second*, to ensure data compatibility, we processed each dataset from scratch using a shared preprocessing pipeline. This standardized procedure topographically aligns subjects to a shared brain-space (the HCP fsLR32k space [43]), thereby mitigating potential confounds arising from differences in experimental parameters like voxel resolution, stimulus duration, and TR [26, 44]. *Third*, to enable robust tests of model generalizability, we carefully curated the aggregated data and stimuli into independent training and testing splits, ensuring that there is no leakage between the two.

**Fig. 1:**
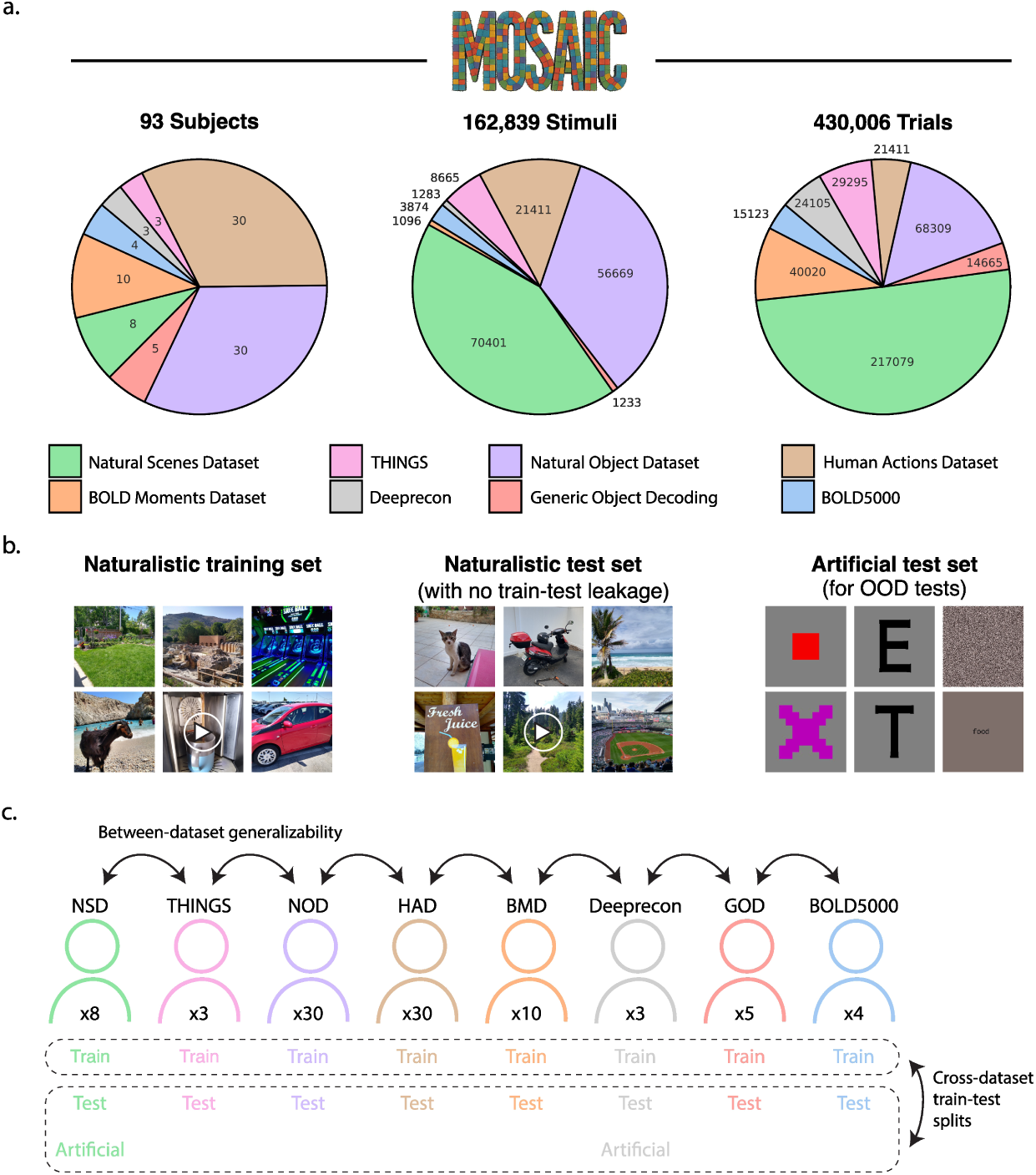
Stimulus set compilation and methodology overview. **a** Pie charts show the subject, stimulus, and single trial contributions of each of the eight large-scale fMRI datasets to the meta dataset after filtering. **b** The aggregated stimulus set is split into a naturalistic train, naturalistic test, and artificial test set. MOSAIC’s naturalistic stimuli span multiple sources, from categorical images to in-the-wild images to short videos. **c** Each subject in MOSAIC has their own train-test split. MOSAIC can be used to run experiments in one subject or dataset and test its generalizability across datasets, or train-test splits can be combined across datasets for data-intensive computational modeling. Representative stimuli for the naturalistic train and test set examples were sourced from the study author, and artificial test set stimuli were sourced from the Natural Scenes Dataset and Deeprecon studies.

The resulting MOSAIC dataset is composed of 162,839 unique images paired with 430,007 single trial fMRI responses. By providing a large-scale, consistently processed, and cleanly partitioned resource, MOSAIC establishes a robust framework for the next generation of data-intensive foundation-level modeling in vision neuroscience. Importantly, preprocessing a dataset within MOSAIC’s framework does not preclude its use for within-dataset analyses. Instead, MOSAIC only adds to its utility by giving other researchers the option to include it in an aggregate set. The following sections detail our stimulus aggregation and fMRI alignment procedures.

#### 2.1.2 Stimulus set aggregation

A primary challenge in aggregating datasets is creating a unified stimulus set with clean training and testing splits. Simply merging the original splits would introduce stimulus duplicates and, more critically, train-test leakage, which would invalidate downstream modeling results. To prevent this, we implemented a rigorous two-stage filtering procedure. As shown in Figure2a, (1) we resolve all direct conflicts where the same stimulus appeared in one dataset’s train set and another’s test set. (2) We used a perceptual similarity threshold (DreamSim [42]) to identify and remove any training images that were highly similar to the images in the final, aggregated test set. We use DreamSim embeddings as this model was optimized to capture human perceptual similarity judgments and shows excellent results on a wide variety of stimuli. The level of similarity that constitutes test set leakage is in part both subjective and dependent on the research question being asked [45], with implementations typically ranging from more strict category-level distinctness [46, 47] to more liberal stimulus-level distinctness [11, 48]. The perceptual similarity exclusion criteria used here compromises between these two implementations. We defined a similarity threshold as the average perceptual similarity of the first and last frames across all 1,102 BOLD Moments Dataset video stimuli, reasoning that if two stimuli were more similar than this threshold, they can be considered perceptual duplicates belonging to the same event as another stimulus (see Supplementary Fig. A1 for examples of excluded stimuli). This process ensures the independence of MOSAIC dataset splits while also maintaining compatibility with the original publications’ proposed splits (see Methods section 7.3 for more details).

#### 2.1.3 fMRI alignment

A wide variety of strategies have been developed to align fMRI data across participants [49–51]. Here we adopt a widely used surface-based approach of registering each subject’s native brain to a shared brain cortical surface template, fsLR32k [52]. This surface-based mapping is widely considered superior to volumetric normalization techniques [53–57] (but see [58]) and enables vertex-to-vertex correspondence across all subjects. Furthermore, the fsLR32k template seamlessly integrates with the Human Connectome Project’s MMP1.0 parcellation [43], which defines 180 regions of interest (ROIs) per hemisphere from 22 larger sections (Fig. 2d). This standardized parcellation simplifies modeling efforts by ensuring ROI data across participants are uniform in size and correspond to similar anatomical locations.

**Fig. 2:**
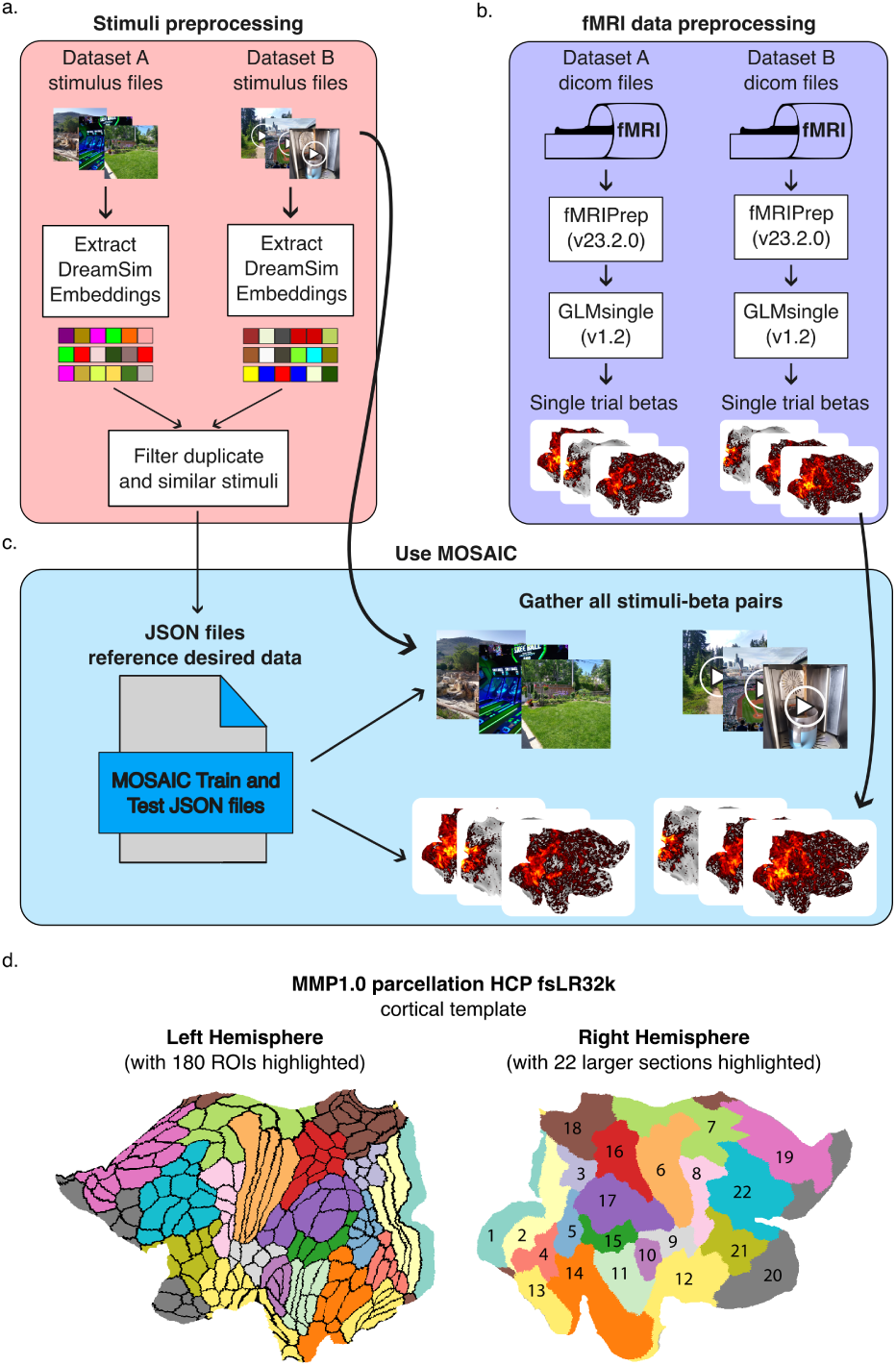
MOSAIC dataset aggregation overview. **a** Combining stimulus sets across datasets (red box). DreamSim [42] features were first extracted for each stimulus. The stimulus filenames and features were then used to resolve conflicts of duplicate and perceptually similar stimuli between datasets to create a naturalistic training, a naturalistic testing, and an artificial testing split that minimize test set leakage. Representative stimuli for the stimulus files examples were sourced from the study author. **b** Combining fMRI data across datasets (purple box). fMRI data were preprocessed using identical fMRIPrep pipelines. The resulting preprocessed fMRI data were then input into a General Linear Model to estimate beta values for each trial. **c** Using MOSAIC (blue box). All stimuli and single trial beta estimates are aggregated, and the curated training and testing split files reference the desired the stimulus-beta pairs. **d** fsLR32k cortical surface mesh and MMP1.0 parcellation. A flattened brain registered to the fsLR32k cortical surface space depicts the MMP1.0 parcellation [43]. The left hemisphere shows the boundaries of the 180 regions of interest (ROIs) within one of 22 color-coded sections. The right hemisphere labels these sections following [43].

We implemented a shared preprocessing pipeline to transform fMRI data from different experimental paradigms (varying TRs, stimulus types, stimulus duration, scanner strength etc.) into a set of consistently formatted fMRI beta-stimulus pairs (Fig. 2b). First, each dataset was processed with an identical configuration of fMRIPrep (v23.2.0) which handled standard preprocessing and registration to the fsLR32k cortical surface [40]. Next, we used GLMsingle to estimate single trial beta values from the preprocessed time series [41]. For datasets with attention-checks highly correlated to stimulus features (i.e. the HAD and NOD datasets), we also regressed out the button press confounds. Finally, to prevent information leakage, the beta estimates of dataset’s train and test splits were normalized using only the mean and standard deviation calculated from the train split betas (see Methods section 7.4 for details).

The file outputs of the stimulus set aggregation step simply reference the desired preprocessed fMRI data for downstream analyses to easily accommodate changes in stimulus set curation or additions to the fMRI data (Fig. 2c).

Towards the goal of supporting data-hungry computational models, MOSAIC additionally makes available all task and resting state (where applicable) fMRI runs in a timeseries format after applying denoising, detrending, and filtering steps after the fMRIPrep preprocessing and fsLR32k space registration (see Methods section 7.4.2) [59, 60]. Task fMRI data in a time-series format may benefit research interested in real-time models of brain activity and latency/accuracy tradeoffs with preprocessing steps [13]. As both MOSAIC and the Human Connectom Project use the fsLR32k registration space, MOSAIC’s resting state brain activity can be readily used with the Human Connectome Project’s hours of resting state data from over 1,000 subjects, which has been previously used to pretrain massive encoder-decoder frameworks that learn powerful latent representations of brain activity [31, 36, 61, 62]. Such methods can also investigate how a MOSAIC subject’s resting state fMRI, when incorporated into the pretraining data, affects a subject’s task-based neural prediction performance.

#### 2.1.4 MOSAIC preprocessing pipeline validation

To validate this pipeline, we compared the vertex-wise noise ceiling signal-to-ratio (NCSNR) reliability metric of the eight NSD subjects computed with beta estimates from MOSAIC’s pipeline in fsLR32k space and beta estimates released by NSD’s pipeline in fsaverage space (see methods section 7.5). In Figure 3a, a visual comparison of subject 01 shows similar spatial patterns and high and low NCSNR between the two pipelines, and the remaining seven subjects show a similar result (see Supplementary Figure A6 for all eight subjects). Figure 3b shows the percent of vertices above a range of NCSNR thresholds between the two pipelines, highlighting that the MOSAIC pipeline (blue) gives comparable results to NSD’s original pipeline (orange) evidenced by their large degree of overlap. Subject 07 shows the least amount of overlap, but its still relatively high reliability and downstream modeling results (see analyses below) confirm that it contains useful signal. We compute and plot NCSNR-based reliability metrics for all subjects and train-test splits or, for HAD and NOD subjects that do not contain stimulus repeats, replicate their validation measures used in their original manuscripts (plots are available alongside the data. See 5). In brief, we find that MOSAIC’s fMRI preprocessing achieves similar or higher reliability than that of the datasets’ original publications likely due to the robust beta estimates from GLMsingle [41].

**Fig. 3:**
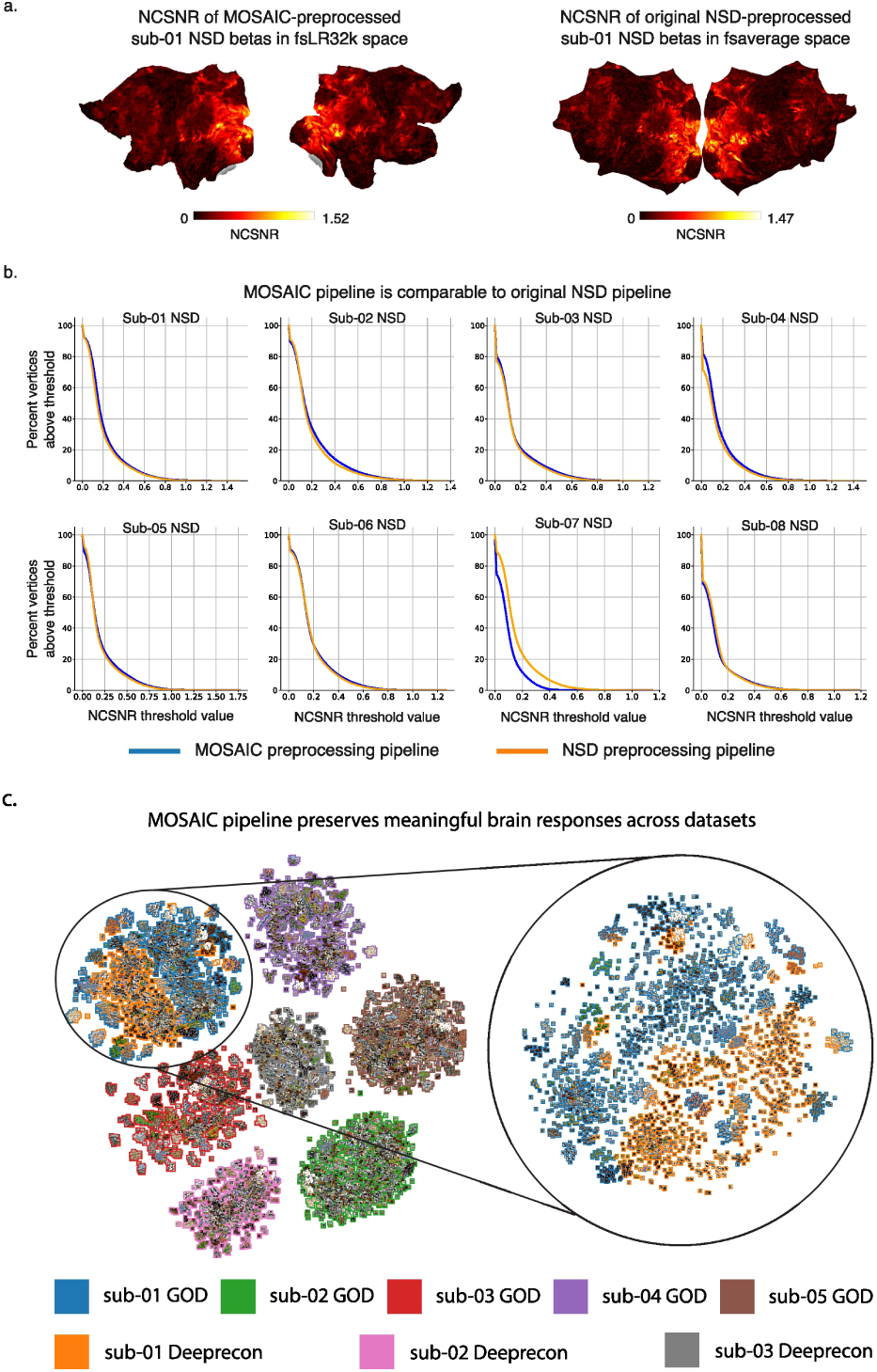
MOSAIC preprocessing pipeline validation. **a** Flat map comparison of noise ceiling signal-to-noise ratio (NCSNR) of Natural Scenes Dataset (NSD) subject 01 between preprocessing pipelines. NCSNR is computed at every vertex from the final beta estimates in MOSAIC’s preprocessing pipeline in fsLR32k space (left) and NSD’s original preprocessing pipeline in fsaverage space (right). Higher NCSNR is better. **b** Percent vertices above NCSNR threshold. For each of the eight NSD subjects, the percent of vertices above a NCSNR value is plotted for both the MOSAIC preprocessed version (blue) and NSD’s originally preprocessed version (orange). **c** Clustering fMRI responses. The TSNE plot (perplexity=20) shows single trial brain responses from visual cortex (MMP1.0 sections 1-5) overlaid with the corresponding image stimulus. The responses from the GOD and Deeprecon datasets primarily cluster around subjects and then stimulus but not dataset. The inset shows a separate TSNE plot focusing on just subject 01 from GOD (blue) and Deeprecon (orange), who is the same individual that participated in both studies. The TSNE plot was initialized from a Multi-Dimensional Scaling dimensionality reduced Representational Dissimilarity Matrix, which was computed from pairwise Pearson correlations between the single trial fMRI responses in visual cortex.

An effective dataset alignment procedure must minimize site-specific technical artifacts from different acquisition sites (e.g., different scanners or location) while preserving meaningful subject-specific biological variation in the brain’s response to stimuli [63]. Does MOSAIC succeed in achieving this balance? To address this question, we leveraged a natural experiment created by the overlap between the Generic Object Decoding (GOD, n=5 subjects) and Deeprecon (n=3 subjects) datasets. Although collected at different sites using different scanners (3T Siemens Trio versus Verio), different brain resolutions (3mm versus 2mm isotropic voxel size), different scanning parameters (3s versus 2s TRs), and at different stimulus durations (9s versus 8s stimulus presentation), these two datasets share 50 naturalistic test stimuli and fortuitously even one individual subject who participated in both studies. This setup allows for a direct assessment of MOSAIC’s validity as an aligned dataset by testing whether the fMRI data for this individual clusters by biological identity rather than by the technical details of the acquisition site.

The results of this natural experiment confirms that MOSAIC’s preprocessing successfully preserves biological identity over technical artifacts. We visualized the single-trial activations in MMP1.0 sections 1-5 for all subjects and stimuli for the GOD (n=35 repetitions per stimulus, 8750 single trials) and Deeprecon (n=24 repetitions per stimulus, 3600 single trials) datasets using TSNE embeddings (Fig. 2c). This visualization reveals two key findings. First, the fMRI responses cluster by individual subject, not by acquisition site. Second, within each subject’s primary cluster, repeated trials of the stimulus form tight sub-clusters, demonstrating high intra-subject reliability. The data from the subject scanned at both sites provides the most compelling evidence of minimal site-specific confounds. Despite being collected on different scanners in different locations, this individual’s data from both studies co-localized in the embedding space, forming a single cluster distinct from all other subjects. This demonstrates that subject-specific neural signatures are far more prominent than any site-specific effects. This result also aligns with prior work showing inter-subject variability to be a much larger source of variance than inter-site variability in multi-site fMRI studies [64–67]. We presume biological identity is a stronger organizing factor than stimulus content in this analysis because the MMP1.0 parcellation used to extract each subject’s activations is not functionally specific and thus contains strong subject-specific patterns [68–72].

To summarize, MOSAIC provides a general and extendable framework for aligning vision fMRI datasets to support a wider range of scientific questions. Crucially, this framework is extensible and not methodologically constrained. New datasets can be incorporated into MOSAIC post-hoc and all data can be re-processed as fMRI preprocessing methods evolve. As we have demonstrated, the resulting fMRI-stimulus pairs successfully minimize site-specific artifacts and retain biologically relevant subject variation. The following sections demonstrate how MOSAIC can be leveraged to replicate known scientific findings across datasets and to train predictive models of the visual system with unprecedented generalizability.

### 2.2 Linear decoding models trained on perceptually diverse stimulus sets are more accurate and less variable

A major goal of cognitive neuroscience is to decode brain responses to determine *what* information is represented in the brain. Linear decoders are a core tool in this effort [13, 45, 47, 73–76], but their performance is naturally constrained by the size and breadth of neural data available for training [77–79]. We used MOSAIC to ask how the composition of the training stimulus set affects a decoding model’s accuracy, stability, and ability to generalize. We leveraged MOSAIC’s scale to systematically vary perceptual diversity and sampling strategy at fixed training size, and evaluate performance on naturalistic in-distribution and artificial out-of-distribution test sets. We hypothesized that perceptually diverse training sets will yield higher accuracy and lower variance, and that boundary-focused sampling will preferentially improve out-of-distribution generalization.

To directly test our hypothesis, we first constructed training sets with maximally different perceptual content for each of the 93 subjects. Using a perceptual feature space defined by DreamSim [42], we sampled two subsets of 200 stimuli: a low-diversity set (the 200 nearest neighbors to a random seed stimulus) and a high-diversity set (iteratively selecting the farthest stimuli starting from a random seed stimulus) (Fig. 4a). We then trained linear ridge regression models on each subset to decode fMRI activity from the visual cortex into this perceptual feature space (Fig. 4b) and evaluated their accuracy on a held-out set of naturalistic images (Fig. 4c). The stimulus sampling and model training process was repeated fifty times with different random stimulus samplings, and results were aggregated over the 93 subjects.

**Fig. 4:**
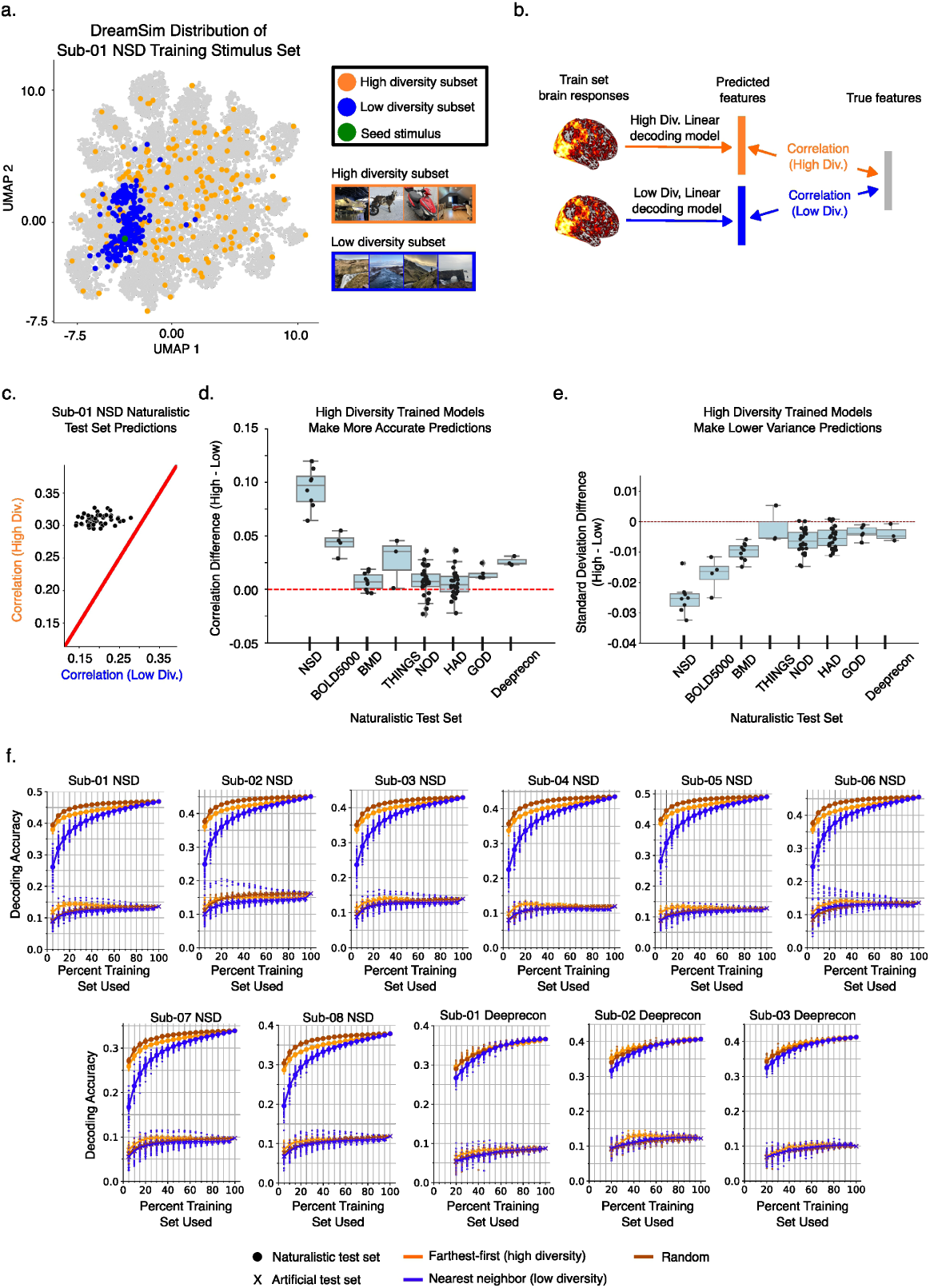
Decoding with high and low diversity stimuli sets. **a** High and low diversity sampling procedure. Out of all stimuli viewed by a given subject, a low (blue) and high (orange) diversity subset was chosen as the closest (nearest neighbors algorithm) and farthest (farthest-first algorithm) 200 stimuli with respect to a random seed stimulus (green). **b** Model training and prediction. Linear decoding models were independently trained to map brain activity from visual cortex (MMP1.0 sections 1-5) to the DreamSim embeddings of the corresponding high and low diversity subsets. The models were then tested to predict DreamSim embeddings of the subjects’ held out naturalistic test sets. Representative stimuli for the high and low diversity stimulus examples were sourced from the study author. **c** Subject 01 NSD prediction results. Representative results from Natural Scenes Dataset subject 01 plot test set predictions from a model trained on high diversity subset and low diversity subset. Decoding accuracy is measured as the Pearson correlation between a stimulus’s predicted DreamSim feature vector and its ground truth feature vector. Each of the fifty points corresponds to a different randomly sampled subset of the subject’s stimulus set. Points above the red diagonal indicate better prediction performance from a model trained with a high diverse subset than one trained with a low diverse subset. **d** High-low diversity prediction accuracy. The mean high minus low difference of the fifty samples per subject are plotted. **e** High-low diversity prediction standard deviation. The high minus low difference in standard deviation of the fifty predictions are plotted. Results in panels **d** and **e** are separated by subject and dataset (black dots) but statistical significance is computed over all subjects (Wilcoxin Signed Rank Test, two-sided against 0, p*<*0.05, n=93). Box plots show the first and third quartiles (large box), median (white dot), and whiskers extending to 1.5 times the interquartile range. Outliers are denoted by diamonds. Individual points represent results of individual subjects. **f** Naturalistic and artificial test set predictions. Linear decoders predict naturalistic (circle, top) and artificial (x, bottom) test sets using nearest neighbor (blue), random (brown), and farthest-first (orange) sampling strategies using increasing amounts of naturalistic training set data. Results from individual subsets are plotted as points.

Models trained on high diversity stimulus subsets had significantly greater prediction accuracy than those trained on low diversity subsets (Fig. 7.7d) (Wilcoxon Signed-rank Test, p*<*0.05, two-sided; n=93, p=6e-10, CLES=0.80). Furthermore, the high stimulus diversity models also exhibited markedly higher stability. They had significantly lower standard deviations in predictions than the models trained on the low diversity stimulus subsets, indicated by the negative high-low standard deviation differences in Fig. 4e (Wilcoxon Signed-rank Test, p*<*0.05, two-sided; n=93, p=8e- 16, CLES=0.92). The long-tailed distributions indicate that stimulus set diversity impacted some subjects and datasets more than others, likely driven by the quality of the corresponding fMRI data.

Building on this finding, we next explored how different sampling strategies interact with training set size and a decoder’s ability to generalize to a new, unseen stimulus distribution. We used the NSD and Deeprecon datasets (which have naturalistic and artificial test set splits) to train decoders on naturalistic images sampled with three strategies: (1) nearest-neighbor (low-diversity, blue) (2) random (brown) (3) and farthest-first (highest-diversity, orange). We tested these decoders on both an in-distribution (naturalistic images) and an out-of-distribution (artificial images) test set 4f). As before, the stimulus sampling and model training process was repeated fifty times, each time with a different random seed. We also varied the number of samples used to train the decoder (to simulate different dataset sizes) from 5% to 100% in 5% increments while ensuring at least 200 training samples (see Methods section 7.7 for details). By design, when 100% of the training samples are used to train the decoder, the three sampling strategies converge to the same result.

For the NSD subjects, the mean predictions of the random sampling strategy consistently outperformed the mean predictions of the farthest-first and nearest neighbor strategies at all training set sizes for the in-distribution naturalistic test set. When predicting the out-of-distribution artificial test set, the farthest-first sampling strategy results in consistently higher mean predictions but only when using up to approximately 50% of the training set. These low-sample mean predictions are often higher than the full training set predictions. The prediction patterns in Deeprecon follow the same trend but were more variable at each training set size. This is likely due to lower fMRI data quality (as measured by vertex reliability) and the fact that their naturalistic test set was designed to not overlap with the train set in terms of semantic categories.

These NSD and Deeprecon results reflect the unsurprising fact that the best sampling strategy is the one most likely to select stimuli with features most similar to the test set. Specifically, the random strategy, compared to the farthest-first strategy, samples throughout the feature space with little concentration at the edges, making it an ideal choice for in-distribution predictions. The farthest-first algorithm samples stimuli with features that lie near the edges of the feature space, meaning it can best select informative samples closest to an out-of-distribution test set. However, too much sampling along these boundaries will inevitably include the most distant, and most irrelevant, samples that can end up hurting decoding performance. We note that the nearest neighbors sampling strategy consistently has the greatest decoding variability and often the worst decoding accuracy in all training set sample sizes, further supporting the conclusions described in Fig. 4d,e. Interestingly, the nearest neighbors strategy can contain high accuracy outliers that outperform the other strategies; in these instances, we qualitatively observe that the random seed and subsequent nearest neighbors happened to identify a cluster of images with features especially relevant for some artificial stimuli, such as a cluster of zebras that likely led to better decoding performance of the black and white spiral gratings found in the NSD artificial test set. Finally, we note that decoding performance for out-of-distribution test sets were significantly worse than the performance for in-distribution sets, emphasizing that while linear decoders can achieve above chance generalization, training on samples with a similar distribution to the testing samples seems necessary for best performance.

These results show that training-set diversity has a measurable impact on decoding accuracy and stability. They also highlight a key difference between MOSAIC, and other large-scale datasets such as the Human Connectome Project [9], UKbiobank [80], Cam-CAN dataset [81], and ABCD dataset [82]: those datasets emphasize subject breadth but offer limited diversity in task stimuli, whereas MOSAIC aggregates richly varied visual stimuli at the trial level. In effect, many fMRI experiments are representative of this low diversity regime: they sample a relatively small stimulus set from a larger dataset defined by a feature space in order to decode that feature space in the brain (e.g., sampling categorically-defined ImageNet or COCO datasets to decode categories). Our results show this low diversity regime consistently leads to lower and more variable decoding accuracy and should encourage researchers to emphasize dataset diversity when sampling future fMRI stimulus sets. Echoing previous work [77–79], we recommend future fMRI datasets first establish a large, highly diverse pool of candidate stimuli from which to sample. As a proxy for sufficiently diverse content, one can attempt to mirror the distribution of recent large-scale computer vision datasets that explicitly target diversity and have successfully been used to train models that exhibit state-of-the-art image or video understanding capabilities [83–85]. With a large and diverse pool of candidate stimuli, random sampling is an effective, feature-agnostic, and simple strategy that can comprehensively cover a distribution. In the event that the distribution of stimulus test sets are not known beforehand, as is often the case, a random sampling strategy can be supplemented with a farthest-first sampling strategy in a feature(s) space that the researcher expects to be relevant in order to include samples along the feature boundary that may be crucial for good generalization.

### 2.3 Brain-optimized encoding models scale across datasets

Beyond decoding *what* information the brain represents (previous section), a second major goal of cognitive neuroscience is to build models that accurately explain *how* sensory inputs are transformed into those neural representations. The last decade has established Deep Neural Networks (DNNs) as the most promising candidates for this endeavor [5, 10, 86–88]. The dominant strategy has been to repurpose ‘task-optimized’ models from the AI community as voxelwise encoding models, which are pre-trained on millions of images for tasks like object classification [45, 73]. An alternative approach is to build ‘brain-optimized’ models by training models end-to-end on neural data itself [12, 89–92], which can offer a more explicit model of sensory processing and incorporate biological constraints starting at model training. However, fully embracing this brain-optimized modeling approach has been challenging because the data requirements of modern DNNs far outstrip the scope of typical neuroimaging experiments. Thus, to successfully train high-parameter, even foundation-level, brain-optimized models, it is necessary to aggregate data across many subjects and datasets and overcome ever-present inter-subject [68–70] and inter-site [67, 93, 94] differences. The MOSAIC dataset is designed to make this feasible by providing a large-scale, aligned collection of trial-level data with curated train-test splits. Using this resource, we compare three architectural frameworks for joint model training and identify a multi-subject architecture that leverages the full scale of MOSAIC to achieve consistently higher prediction performance than its traditional, single-subject counterparts.

We use a DNN architecture following [12] with a shared core encoder network followed by a factorized linear readout (Fig. 5a) [95]. The linear readout constrains all nonlinear computations to the core, which improves systems-level interpretability and hypothesis testing. The readout head also contains drastically fewer parameters than fully connected layers, allowing the system to easily scale to multiple subject-specific readout heads and high dimensional output predictions. This brain-optimized framework also makes multivariate predictions of brain activity, in contrast to the more commonly used voxelwise encoding models that predict each vertex independently from one another [45, 73, 89].

**Fig. 5:**
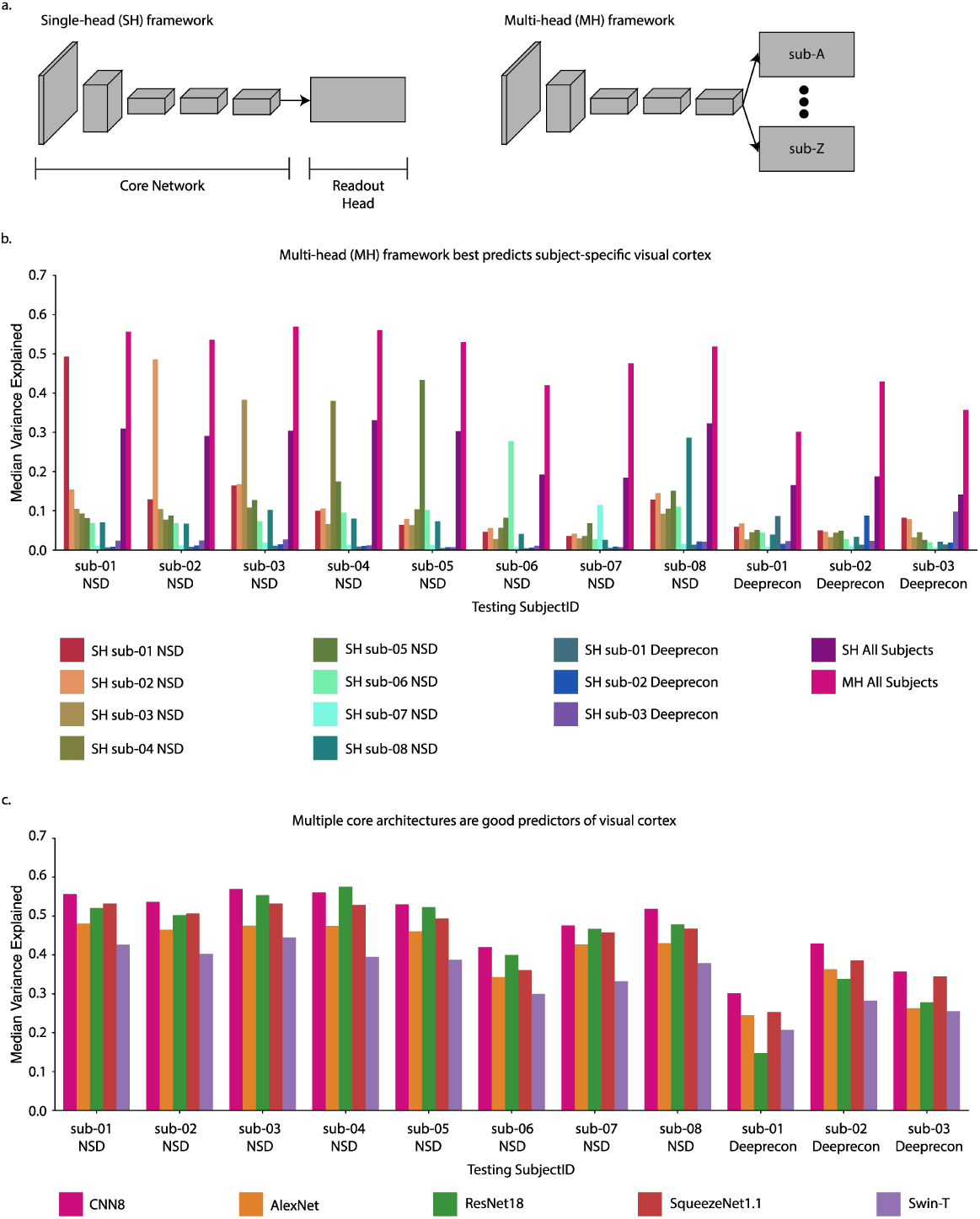
Brain-optimized networks are accurate predictors of visual cortex. **a** Generic brain-optimized network frameworks. A brain-optimized network consists of a core network and at least one linear readout head to map stimulus input into a vector of brain activity. Multiple readout heads can be attached to the core network for subject-specific brain activity prediction. **b** Brain-optimized framework evaluation. Single-head (SH) single subject, single-head (SH) all 93 subjects (deep purple), and multi-head (MH) all 93 subjects (magenta) models predict the test set responses of Natural Scenes Dataset (NSD) subjects 01-08 and Deeprecon subjects 01-03. Prediction accuracy is the voxelwise correlation between the subjects’ predicted and true brain responses within visual cortex vertices (MMP1.0 sections 1-5). The median of each vertex’s squared correlation divided by its noise ceiling, or explained variance, is plotted. **c** Prediction performance of brain-optimized models using different core network architectures. Five multi-head brain-optimized models using different core networks were trained to predict visual cortex responses (MMP1.0 sections 1-5). The 8 block CNN (magenta), AlexNet (orange), ResNet18 (green), SqueezeNet1.1 (maroon), and Swin-T (purple) core networks vary in parameter count, task accuracy, and architecture family (e.g., CNN and transformer) yet all are good predictors of visual cortex.

We tested three brain-optimized frameworks to identify the most accurate method for predicting subject-specific fMRI activity in visual cortex using MOSAIC. Each framework uses the same eight layer convolutional core (CNN8, following [12]), but they differ in their training data and readout architecture. The three frameworks are as follows: (1) The *baseline model* is a standard single-head (SH) model trained using data one subject at a time. This model circumvents inter-subject and inter-site differences and, for subjects from the Natural Scenes Dataset (NSD), reflects performance achievable with extensive single-subject data collection approaching practical limits [14]. (2) The *pooled model* is a simple pooling strategy with a single-head (SH) trained on the combined data from all 93 MOSAIC subjects. This one-size-fits-all model shares all parameters, forcing it to learn a single, subject-agnostic representation. (3) The *multi-head model* implements a hybrid approach. This model features a shared core network trained on all 93 subjects but routes its output to 93 separate, subject-specific readout heads. This architecture allows the model to learn general visual features from the entire dataset while tailoring predictions to each individual’s unique neural topography. All models are trained from randomly initialized weights. We show prediction results on NSD subjects 01-08 and Deeprecon subjects 01-03 (see Methods section 7.9 for model training information).

Our results indicate that the multi-head model (magenta) was the overall superior modeling framework for MOSAIC. This framework successfully leveraged the shared core network when trained on MOSAIC’s diverse stimulus set across all 93 subjects, and consistently achieved better predictions of visual cortex than the single subject models (Fig. 5b) (see Supplementary Fig. A8 for raw correlation results from the multi-head model on all 93 subjects and test sets). This performance gain appeared to be driven primarily by more accurate predictions in higher-level visual cortex (see Supplementary Fig. A4). We also observed an interesting trade-off with the pooled model (deep purple); while it underperformed for subjects with high-quality data, it was more accurate than the single-subject models in low-data or low-quality settings, suggesting that training on a group of subjects can capture highly predictive but subject-agnostic representations of vision [96] (see Supplementary Fig. A3 for noise ceiling comparison across subjects). Finally, the baseline single-head models trained on one only one subject and tested on a held-out individual unsurprisingly yielded the worst performance (Fig. 5b). This highlights that some subject-specific parameters are probably essential for accurate predictions.

We next confirm that the multi-head brain-optimization framework is robust and generalizes well to other models (different architectures, varying parameters and task capabilities). We found that the eight-layer CNN (CNN8, used above), AlexNet [97], SqueezeNet1.1 [98], ResNet18 [99], and Swin-T (i.e., Swin Tiny) [100] vision transformer all achieved strong brain prediction performance (Fig. 5d) (see Supplementary Table A1 for core network comparisons and Methods section 7.8 for descriptions of the architectures). In general, the four convolutional networks typically outperformed the Swin-T vision transformer network at predicting brain responses even though Swin-T can be optimized to obtain a notably higher performance task itself. SqueezeNet1.1 matched or slightly exceeded AlexNet at predicting responses despite using less that one-third of the parameters in the core network. Although ResNet18 implemented brain-like recurrent computations [101–104], it was not obviously more predictive than other models. We further demonstrate that these brain-optimized models can be used in a voxelwise encoding paradigm on a non-MOSAIC fMRI dataset [105], where features from the brain-optimized models largely outperformed their task-optimized counterparts in predicting high-level brain responses in an external fMRI dataset (Supplementary Fig. A7a,b)

These results, made possible by the scale and diversity of MOSAIC, demonstrate that the multi-head architecture is a viable path for scaling brain-optimized models. By jointly training on multiple subjects and datasets, we can achieve better prediction performance than what is possible with using only extensive single-subject data. This joint training is crucial, as it forces the model to learn a shared core that reflects robust, generalizable properties of vision [34]. Training ablations show that a model trained on only the eight NSD subjects achieves better prediction accuracy than the model trained on all MOSAIC (Supplementary Fig. A5b) but, as suggested by Fig. 5b, this performance increase on NSD subjects comes at the expense of poor generalization to the remaining held-out subjects. Additionally, training the model on one dataset and fine tuning subject-specific readouts would prevent each subject and dataset from contributing to the shared core. Looking forward, a key challenge will be to develop training paradigms where the addition of new subjects or datasets—even noisy ones—can only improve, and never harm, overall model performance.

### 2.4 Brain-optimized models recapitulate subject-specific functional contrasts

Given the brain-optimized models’ excellent prediction accuracy, we next ask if these models are scientifically useful by performing an in silico functional localizer experiment to predict functional regions of interest (fROIs). We first train a new multi-head model to predict the whole brain in order to more closely resemble the data that would be collected in a real fMRI experiment. We train the model on NSD subjects 01-08 instead of all 93 subjects due to GPU memory constraints. We then run subject-specific model inference on a localizer stimulus set (no overlap with the MOSAIC training set) and perform the localizer’s prescribed contrasts to identify face-, place-, body-, and word-selective regions (see Methods section 7.10 for more details) [106]. We note that the image statistics of the localizer’s grayscale, isolated object stimuli on a textured background are out-of-distribution with respect to the model’s primarily colored naturalistic training set. We compare our model’s predicted contrasts with each subject’s category-selective regions independently defined and preprocessed in the original Natural Scenes Dataset experiment using the same localizer task [106]. We regard these experimentally defined categorical contrasts as the ground truth.

All eight subjects predict their four functionally selective regions well above chance levels, as shown in the precision-recall curves (Fig. 6a) and Table 1. We then threshold the predicted contrasts at the value that maximizes the F1 score for each subject and contrast to obtain a binary prediction of the category-selective regions. Manual examination of the top 100 experimental stimuli with the highest mean activation across these predicted category-selective regions reveal clear category membership (see Supplementary Figs. A9-A16 for the stimuli). Visual comparisons between the subject-aggregated predictions and ground truth show that despite no spatial or contiguity constraints were enforced, the predicted regions were often tightly grouped in visual cortex, largely matching the ground truth (Fig. 6b). The word-selective region predictions, while still well-above chance level, were noticeably worse across all subjects. The stimuli that most strongly activate the predicted word-selective regions contain a mix of characters, as expected, but also high frequency content and character-like naturalistic images (e.g., a circular doughnut), suggesting that the model is capturing more of the statistical structure of characters and less the specialized function of word-selectivity.

**Fig. 6:**
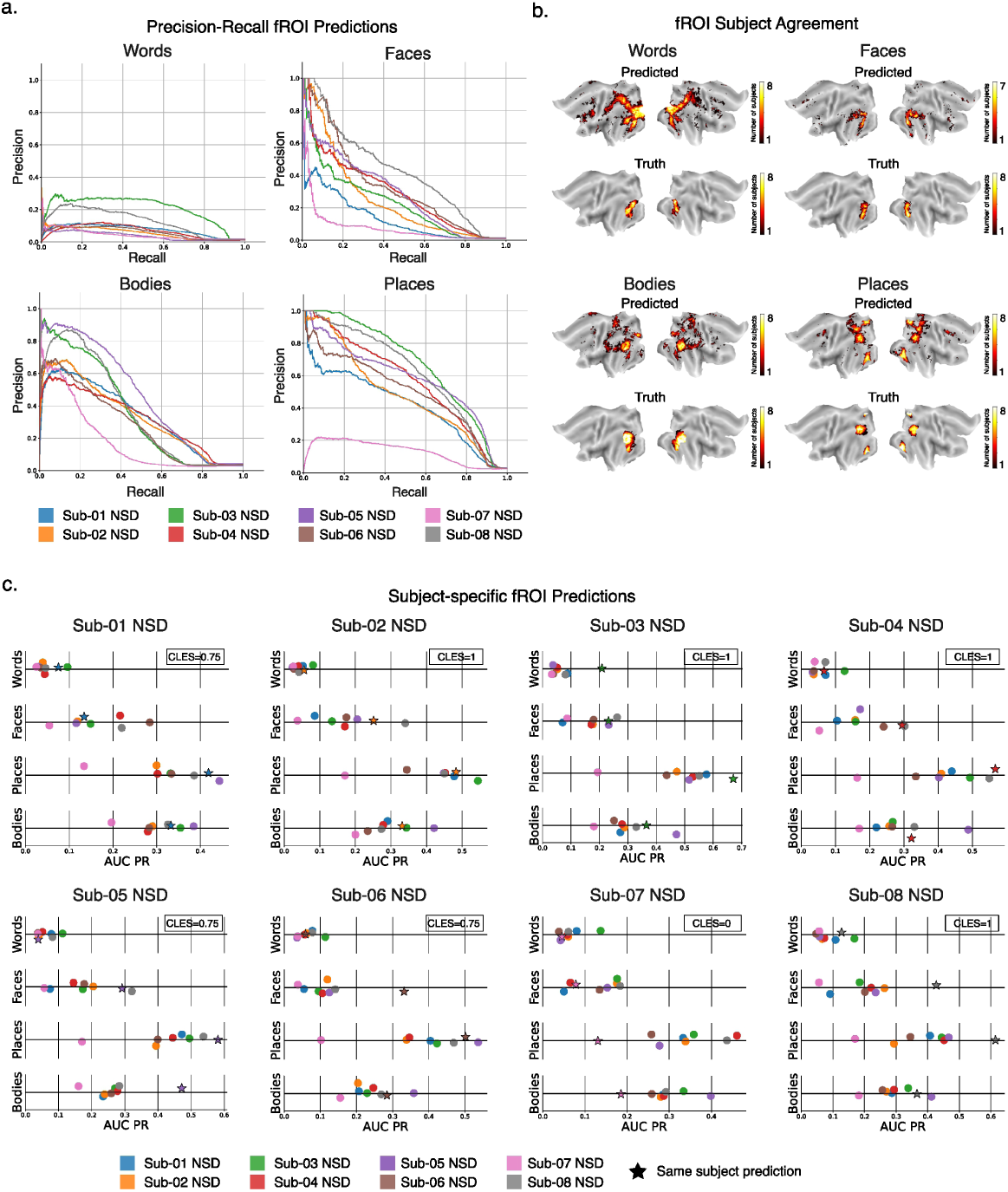
Brain-optimized models for functional ROI prediction (fROI). **a** Precision-recall curve of categorical contrasts. For each of the four word, face, place, and body contrasts, eight precision-recall curves are constructed to measure how well each subject-specific model prediction overlaps with that subject’s ground truth contrast. **b** fROI subject agreement. The contrast value that achieves the greatest F1 score for each subject and contrast is used to threshold the contrast into a binary prediction. The union of each subject’s binary predictions are compared to the union of each subject’s binary ground truth fROIs on a flatmap. **c** Measuring the subject-specificity of fROI predictions. The area under the precision recall curve (AUC PR) for each subject’s synthetic fROI prediction compared with each subject’s ground truth is plotted on a number line. The star represents that the same subject is used for both the prediction and ground truth comparison.

**Table 1:** Area under precision recall curve for functional contrast prediction. . A precision-recall curve is computed for each subject-specific contrast prediction compared to that subject’s ground truth contrast. The area under precision recall curve metric (AUC PR) summarizes the prediction accuracy. The random baseline is the performance of a random binary classifier and is equivalent to *p/*(*p* + *n*), where p is the number of positive examples (vertices in the ROI) and n is the number of negative examples (vertices not in the ROI).

| Subject | Contrast |  |  |  |  |  |  |  |
| --- | --- | --- | --- | --- | --- | --- | --- | --- |
|  | Words (AUC PR) |  | Bodies (AUC PR) |  | Faces (AUC PR) |  | Places (AUC PR) |  |
|  | Prediction | Random | Prediction | Random | Prediction | Random | Prediction | Random |
| Sub-01 NSD | 0.075 | 0.015 | 0.332 | 0.029 | 0.134 | 0.009 | 0.418 | 0.027 |
| Sub-02 NSD | 0.056 | 0.013 | 0.332 | 0.031 | 0.251 | 0.012 | 0.483 | 0.027 |
| Sub-03 NSD | 0.209 | 0.012 | 0.365 | 0.033 | 0.232 | 0.012 | 0.670 | 0.029 |
| Sub-04 NSD | 0.066 | 0.011 | 0.322 | 0.036 | 0.295 | 0.013 | 0.567 | 0.027 |
| Sub-05 NSD | 0.039 | 0.012 | 0.472 | 0.043 | 0.292 | 0.015 | 0.581 | 0.029 |
| Sub-06 NSD | 0.059 | 0.014 | 0.284 | 0.038 | 0.332 | 0.009 | 0.501 | 0.025 |
| Sub-07 NSD | 0.045 | 0.019 | 0.186 | 0.030 | 0.078 | 0.008 | 0.131 | 0.024 |
| Sub-08 NSD | 0.127 | 0.015 | 0.366 | 0.034 | 0.427 | 0.014 | 0.614 | 0.028 |

Next we evaluate the subject-specificity of the model by comparing the Area under Precision Recall Curve (AUC PR) between all subject-specific predictions and one subject’s ground truth (Fig. 6c). If the model’s predictions are subject-specific, the same-subject AUC PR score (denoted by a star) should rank higher (farther to the right on the number line) than the other-subject AUC PR scores. In general, we observe a pattern of same-subject predictions ranking higher than other-subject predictions (except subject 07) and quantify this by computing the effect size of the same-subject rank against the other-subject average rank across the four conditions (CLES=0.75, 1.0, 1.0, 1.0, 0.75, 0.75, 0, 1 for subjects 01-08, respectively). Due to low sample size (n=4 contrasts) and multiple comparisons (8 subjects), statistical tests against a p-value of 0.05 or lower are not appropriate but more extensive investigations are warranted from the large effect sizes. Because fROIs are expected to significantly overlap between subjects [71, 72], one subject’s good fROI prediction will score highly against other all other subjects’ ground truth, often overshadowing the same-subject score (e.g., Subject 03 for words, Subjects 06 and 08 for faces, subject 05 for bodies show strong performance across all subjects).

Taken together, this brain-optimized encoding model can effectively recover subject-specific category-selective regions, thereby demonstrating that these brain-optimized models achieve not just high encoding accuracies but are also scientifically useful. When paired with algorithmic fROI defining methods like Group-constrained Specific-Subject [71], these models may offer a fully automated pipeline for subject-specific ROI identification. This multi-head framework was able to scale its prediction output from an already large 7,831 vertices (visual cortex, MMP1.0 sections 1-5) to 57,051 vertices (whole brain, all MMP1.0 sections) with only minor loss in visual cortex prediction performance (Supplementary Fig. A5a). Even still, figuring out how to scale the output prediction size with no loss in performance will be a crucial step for investigating whole-brain functional connections.

## 3 Conclusion

With MOSAIC, individual fMRI datasets can seamlessly contribute to replicable, generalizable, and data-intensive investigations into human vision. MOSAIC’s scale can be leveraged in two ways - first, hypotheses can be independently tested across multiple datasets to see if results generalize (Fig. 4), and second, fMRI-stimulus pairs can be aggregated across datasets for large-scale model training (Fig. 5). As an example of the former approach, we show how linear decoding models trained on perceptually diverse stimuli result in better and more stable predictions. Following the second approach, we train brain-optimized models across 93 subjects to learn a robust mapping between stimulus and fMRI response. MOSAIC’s already large scale and ability to accommodate new datasets offers a strong foundation to build and evaluate robust models of vision.

MOSAIC uses a shared, state-of-the-art preprocessing pipeline to minimize undesired variability that can arise from seemingly minor differences in software versions, let alone full pipelines. Relatedly, this shared preprocessing pipeline can support investigations into subject- and dataset-specific sources of variability, potentially leading to improved preprocessing methods and a deeper understanding of the sources of inter-subject differences [25]. In line with this effort, the MOSAIC dataset compilation includes edge cases of 16 shared subjects, thousands of shared stimuli between 2 or more datasets, and two datasets (GOD and Deeprecon) with identical stimulus sets, one common subject, but different experimental protocols. In Fig. 2c we leverage these similarities and differences between GOD and Deeprecon to show that MOSAIC’s shared preprocessing pipeline is able preserve biologically meaningful signal while reducing inter-dataset sources of variability.

Importantly, aggregating individual fMRI datasets naturally diversifies the compiled stimulus set. Stimuli sampled from a single source often include biases that undesirably affect model performance [107–109] and are limited in their ecological validity. In aggregate, however, MOSAIC’s naturalistic stimulus set samples six computer vision datasets that include images, short videos, focused categorical content, and in-the-wild contexts that are crucial for evoking a representative range of neural responses [18, 110–112]. Furthermore, we expand on previous work about the importance of a diverse stimulus set, finding that a perceptually diverse stimulus set leads to more accurate and stable brain-to-feature decoding performance (Fig. 4) and compare stimulus sampling strategies across different sample sizes. These results carry direct implications for how future fMRI datasets should sample their stimulus set for the purpose of in- or out-of-distribution decoding; random sampling from a large pool of stimuli achieves excellent in-distribution performance, and farthest-first sampling with respect to a target feature space can aid in out-of-distribution decoding. MOSAIC’s extensible framework is uniquely positioned to allow researchers to explore decoding of new feature spaces as datasets can be added post-hoc to fill gaps in stimulus diversity. Increasing dataset scale has enabled landmark discoveries in computer vision and computational neuroscience, such as describing emergent properties of vision models [113–116], detailing the effects of model architecture, training diet, and objective function on brain-alignment [10, 117–121], and measuring the convergence of multi-modal model representations at scale [122]. Extending these efforts in increasingly large-scale brain-optimized models would help anchor these seminal findings within neuroscience, as such models are explicitly formulated to capture neural sensory processing [89–91, 95]. MOSAIC provides a powerful framework for overcoming the severe limitations of current fMRI dataset scale. In this work we take the first step in overcoming these challenges by showcasing a suite of highly predictive brain-optimized models that can effectively scale across subjects, datasets, and output sizes (Fig. 5). Their multivariate prediction directly to cortical vertices opens new avenues for studying the model’s sub-networks and units that give rise to functional cortical activity. With confidence from their ability to recover subject-specific functional regions (Fig. 6), MOSAIC-trained models can be used to synthesize neural responses on the order of billions of stimulus-brain activity pairs to achieve a dataset size on par with modern artificial neural network vision models [123]. Taken together, MOSAIC serves as a community-led data foundation to enable the next generation of vision neuroscience discoveries.

## 4 Limitations

MOSAIC’s shared preprocessing pipeline between datasets is intended to be lightweight, transparent, and flexible to facilitate the addition of future fMRI datasets and seamlessly adapt to future methodological improvements. Custom preprocessing pipelines might optimize dataset quality on a per-dataset basis. Relatedly, the pre-processing pipeline resamples all fMRI data to approximately 2mm spacing on the fsLR32k cortical surface and thus does not take advantage of some datasets’ higher spatial resolution.

Aggregating independent datasets naturally introduces stimuli and data quality imbalances. This version of MOSAIC is dominated by the Natural Object Dataset (*>* 57,000 image stimuli) and notably high quality Natural Scenes Dataset (*>* 70,000 image stimuli), thus potentially biasing results following the experimental design choices of those study authors. Multi-subject and multi-dataset analyses may also obscure otherwise statistically significant results due to inter-subject and inter-dataset variation. However, results that do reach statistical significance are more likely to be true as they would need to overcome these additional sources of variation.

In-domain data samples must be defined in terms of both (1) the stimulus and (2) the cognitive process. (1) This compilation of fMRI datasets, while large, still does not sample the scale and diversity of a human’s multimodal and embodied visual diet. (2) Moreover, these fMRI experiments measure brain activity during passive viewing of visual stimuli and thus do not achieve a representative diversity of neural processes (e.g., evoked emotions, memory recall of subjective experiences, etc.).

## 5 Data and code availability

MOSAIC fMRI data and model weights are available here:

1. https://aws.amazon.com/marketplace/pp/prodview-vsoockzeptxzw

MOSAIC preprocessing code is available here:

1. https://github.com/blahner/mosaic-preprocessing

Code supporting manuscript results will be released at time of peer-reviewed publication.

Original fMRI datasets and their respective stimuli can be accessed here:

1. BOLD5000: https://openneuro.org/datasets/ds001499
2. BOLD Moments Dataset (BMD): https://openneuro.org/datasets/ds005165
3. Generic Object Decoding (GOD): https://openneuro.org/datasets/ds001246
4. Deeprecon: https://openneuro.org/datasets/ds001506
5. Human Actions Dataset (HAD): https://openneuro.org/datasets/ds004488
6. Natural Object Dataset (NOD): https://openneuro.org/datasets/ds004496
7. Natural Scenes Dataset (NSD): https://registry.opendata.aws/nsd/
8. THINGS: https://openneuro.org/datasets/ds004192
9. functional localizer experiment: https://github.com/VPNL/fLoc

The MOSAIC dataset (preprocessed fMRI data and test-train splits), model weights, and code will be available upon publication in a peer-reviewed journal. Note that the MOSAIC dataset is (1) the eight fMRI datasets preprocessed with the same preprocessing pipeline and (2) a curated test-train split across datasets to control for test set leakage. Reproducing the MOSAIC dataset involves downloading the raw Nifti data from each of the eight original dataset’s publication (links above) and passing them through MOSAIC’s preprocessing pipeline.

## 6 Acknowledgements

We thank Kendrick Kay for giving us early access to the NSD synthetic dataset. Thank you to the Amazon Open Data Sponsorship Program for supporting the dissemination of MOSAIC.

This research was funded by the Multidisciplinary University Research Initiative (MURI) award by the Army Research Office (grant No. W911NF-23-1-0277) to A.O.; the Pathway to Independence Award by the National Institute of Health (NIH) (award No. R00EY032603) to N.A.R.M.

## Declarations

The authors declare no conflicts of interest.

## 7 Methods

### 7.1 Participants

The MOSAIC framework comprises 8 open-access datasets collectively encompassing 93 subjects (Deeprecon=3; THINGS-fMRI (THINGS)=3; BOLD5000=4; Generic Object Decoding (GOD)=5; Natural Scenes Dataset (NSD)=8; BOLD Moments Dataset (BMD)=10; Human Actions Dataset (HAD)=30; Natural Object Dataset (NOD)=30) viewing images or short videos. Sixteen subjects participated in two studies, as determined by correspondence with the study authors (one between GOD and Deeprecon and 15 between NOD and HAD). Unless stated, we treat these 16 individuals as 32 unique subjects to account for potential experiment-specific differences and time-dependent changes in brain responses that may have influenced brain responses. Together, the aggregated MOSAIC dataset thus contains 77 unique human individuals (29 male, 48 female).

### 7.2 Individual dataset experimental designs and stimulus sets

We summarize the relevant details of each dataset’s experimental design and stimulus set in the text below and in Tables 2 and 3. Some datasets may have performed additional tasks (e.g., retinotopy, localizer, venograms etc.) that we do not describe here. We describe the process for filtering out duplicate or highly similar stimuli across datasets in a subsequent section. For additional details, see the dataset’s original publication.

**Table 2:** Stimulus sets and acquired trials of the datasets that comprise MOSAIC. 2,853 filenames of the aggregated stimuli entries across the datasets were non-unique due to 2,805 stimuli being repeated across two datasets and 24 stimuli being repeated across 3 datasets (2,829 overlapping stimuli, 2,853 non-unique filename entries). The 2,853 non-unique filename entries are attributed to 1,751 entries only in the training set (from 1,729 stimuli), 70 entries only in the testing set (from 68 stimuli), and 1,032 entries mixed between the training and testing sets (from 1,032 stimuli). Of the 1,032 mixed entries, the subsequent test/train filtering step removed 1,008 entries from the training set and 24 entries from the testing set. The similarity filter, by design, only removed stimuli from the training set. * Note that in this study we treat all 93 participants (despite the 16 duplicate participants) as different to account for possible experiment-specific and changes within a subject across time that may influence their measured brain responses.

| Dataset | Num. Sub-jects | Num. Stim-uli | Num. Nat. Train Stim. | Num. Nat. Test Stim. | Num. Arti-ficial Test Stim. | Num. Trials | Num. Nat. Train Trials | Num. Nat. Test Trials | Num. Arti-ficial Test Trials |
| --- | --- | --- | --- | --- | --- | --- | --- | --- | --- |
| BOLD5000 | 4 | 4916 | 4803 | 113 | 0 | 18870 | 17255 | 1615 | 0 |
| BMD | 10 | 1102 | 1000 | 102 | 0 | 40200 | 30000 | 10200 | 0 |
| GOD | 5 | 1250 | 1200 | 50 | 0 | 14750 | 6000 | 8750 | 0 |
| Deeprecon | 3 | 1300 | 1200 | 50 | 50 | 24360 | 18000 | 3600 | 2760 |
| HAD | 30 | 21600 | 16200 | 5400 | 0 | 21600 | 16200 | 5400 | 0 |
| THINGS | 3 | 8740 | 8640 | 100 | 0 | 29520 | 25920 | 3600 | 0 |
| NOD | 30 | 57120 | 45696 | 11424 | 0 | 68760 | 55008 | 13752 | 0 |
| NSD | 8 | 70850 | 69566 | 1000 | 284 | 218312 | 191882 | 21118 | 5312 |
| Aggregate | 93 | 166878 | 148305 | 18573 | 334 | 436372 | 360265 | 68035 | 8072 |
| Overlap | (16) | (2853) | (1751) | (70) | 0 | 0 | 0 | 0 | 0 |
| Test/Train Filter | 0 | 0 | (1008) | (24) | 0 | (3607) | (3583) | (24) | 0 |
| Similarity Filter | 0 | (1186) | (1186) | 0 | 0 | (2758) | (2758) | 0 | 0 |
| **Total** | **77\*** | **162839** | **144360** | **18145** | **334** | **430007** | **353924** | **68011** | **8072** |

**Table 3:** Experimental protocols of the datasets that compose MOSAIC. + In GOD and Deeprecon, the images were flashed at a frequency of 2Hz for the duration of their presentation. * In NOD, the ImageNet images were presented for a duration of 1 second and the COCO images were presented for a duration of 0.5 seconds.

| Dataset | TR (sec) | Scanner Strength (Tesla) | Functional Voxel Resolution (mm) | Stimulus Duration (sec) | Visual Angle (deg x deg) |
| --- | --- | --- | --- | --- | --- |
| BOLD5000 | 2 | 3 | 2x2x2 | 1 | 4.6 |
| BMD | 1.75 | 3 | 2.5x2.5x2.5 | 3 | 5 |
| GOD | 3 | 3 | 3x3x3 | 9<sup>+</sup> | 12 |
| Deeprecon | 2 | 3 | 2x2x2 | 8<sup>+</sup> | 12 |
| HAD | 2 | 3 | 2x2x2 | 2 | 16 |
| THINGS | 1.5 | 3 | 2x2x2 | 0.5 | 10 |
| NOD | 2 | 3 | 2x2x2 | 0.5, 1* | 16 |
| NSD | 1.6 | 7 | 1.8x1.8x1.8 | 3 | 8.4 |

#### 7.2.1 BOLD5000

The BOLD5000 dataset [124] was collected in Pittsburgh, PA USA. Subjects 1-3 completed 15 core experimental sessions and subject 4 completed 9 core experimental sessions of naturalistic image viewing. Images assigned to the training and testing splits were presented in random order and could appear in the same run. Images were presented at 4.6 degrees of visual angle for a duration of 1 second followed by a fixation period of 9 seconds. Task vigilance was assessed by asking subjects to press a button with their right hand (all subjects were right-hand dominant) to record their valence judgments of ”like”, ”neutral”, or ”dislike” for each stimulus. Button presses were recorded during the fixation period.

BOLD5000 includes 4,916 naturalistic images (n) sampled from the COCO [125] (n=2,000), the Scene Understanding (SUN) database [126] (n=1,000), and the ImageNet dataset [127] (n=1,916). The training set contains 4,803 images, each presented once per subject. The testing set contains 113 images, proportionally representative of the three databases (2/5 COCO, 1/5 SUN, 2/5 ImageNet), presented 1–5 times per subject.

#### 7.2.2 BOLD Moments Dataset

The BOLD Moments Dataset [15] was collected in Cambridge, MA USA. All ten subjects completed 4 sessions of the main experiment. Testing set and training set videos were presented in their respective testing and training runs. A trial consisted of the 3 second video presentation at 5 degrees of visual angle overlaid with a center fixation cross followed by a 1 second fixation period. Subjects were instructed to press a button during blank trials (no video was presented), during which the fixation cross also dimmed color.

BMD includes 1,102 3-second videos sourced from the Multi-Moments In Time dataset [128], a subset of the Moments in Time dataset [129]. The stimulus set was split into a training set of 1,000 videos (presented 3 times per subject) and a testing set of 102 videos (presented 10 times per subject). All subjects viewed all stimuli the same number of times. The videos were shot by amateur videographers (e.g., ’home videos’) and depict animate and inanimate naturalistic content.

Each subject also underwent 40 minutes of resting-state fMRI scans. Subjects were instructed to keep their eyes closed, not think of anything specific, and remain awake.

#### 7.2.3 Generic Object Decoding

The Generic Object Decoding (GOD) [46] dataset was collected in Kyoto, Japan. Each of the 5 subjects viewed naturalistic training images across 3 sessions and naturalistic testing images across 4 sessions. A one-back vigilance task was used during the naturalistic image presentations to measure the subject’s attentiveness. The naturalistic images were presented at 12 degrees of visual angle for a duration of 9 seconds flashed at 2Hz with an overlaid center fixation. No rest block followed the image presentation.

The GOD dataset contains 1,200 naturalistic training images (8 images each from 150 distinct ImageNet categories) and 50 naturalistic testing images (1 image from each of the 50 distinct non-overlapping ImageNet categories). Training images were presented once per subject, while testing images were presented 35 times per subject.

#### 7.2.4 Deeprecon

The Deeprecon dataset [47] was collected in Kyoto, Japan, and presented naturalistic training images across 15 sessions, naturalistic testing images across 3 sessions, artificial shape images across 2-3 sessions, and artificial letter images across 1 session for each of the 3 subjects. The authors report high accuracy on a one-back button press vigilance task. Stimuli were presented with a central fixation for a duration of 8 seconds flashed at 2Hz subtending 12x12 degrees of visual angle. The artificial letter stimuli were followed by a 12 second rest block. There were no rest blocks for the naturalistic train, naturalistic test, and artificial shape images.

The Deeprecon dataset includes 1,200 naturalistic training images and 50 naturalistic testing images from the ImageNet database [127], identical to those used in the GOD dataset. Additionally, it contains 50 artificial testing images (10 letters and 40 shapes). Each naturalistic training image, naturalistic testing image, artificial shape image, and artificial letter image was presented 5, 24, 20, and 12 times, respectively, per subject.

#### 7.2.5 Human Actions Dataset

The Human Actions Dataset (HAD) [130] was collected in Beijing, China. Each of the 30 subjects participated in 1 session where they viewed 720 unique video stimuli. Each trial consisted of the 2 second video presentation followed by a 2 second fixation period. The videos subtended 16 degrees of visual angle overlaid with a central dot fixation. Subjects were instructed to press one of two buttons with their right or left thumb during the 2 second inter-stimulus fixation period to indicate if the recently viewed video contained a ”sport” or ”non-sport” action, respectively.

HAD includes 21,600 2-second videos from the Human Action Clips and Segments (HACS) dataset [131]. Each video contains a naturalistic human action and belongs to one of 180 action categories. Each subject viewed 720 unique videos (4 per action category), with no videos repeated within or across subjects. The authors did not specify a training or testing split; however, for the purpose of MOSAIC, we defined a test set as the videos presented during each subject’s final 3 of 12 runs (3 runs x 60 videos/run = 180 videos, one per action category). The remaining 540 videos per subject were designated as the training set. This approach ensures consistent representation across all action categories in both sets.

#### 7.2.6 Natural Object Dataset

The Natural Object Dataset (NOD) [132] was collected in Beijing, China. Subjects 1-9 participated in 4 ImageNet sessions and 1 COCO session. Subjects 10-30 participated in 1 ImageNet session and zero COCO sessions. During each ImageNet session, subjects viewed 1,000 unique ImageNet images presented with a duration of 1 second at a 16 degree visual angle overlaid with a fixation dot followed by a fixation period of 3 seconds. Subjects were instructed to press a button during the 3 second fixation period with their right thumb if the image depicted an animate object and their left thumb if the image depicted an inanimate object. During a COCO session, images were presented with a duration of 0.5 seconds at a 16 degree visual angle overlaid with a fixation dot followed by a fixation period of 2.5 seconds. Subjects were instructed to press a button when they observed the fixation dot change color, which occurred with 50% probability only on blank trials.

NOD has 57,120 unique naturalistic images from the ImageNet [127] (n=57,000) and COCO [125] (n=120) databases. Subjects 1-9 viewed 4,000 ImageNet images one time and 120 COCO images 10-11 times. Subjects 10-30 viewed 1,000 ImageNet images one time. No ImageNet images were repeated within or across subjects, and the 120 COCO images were shared between the first nine subjects. Each ImageNet session showed 1,000 ImageNet images (4 ImageNet sessions for subjects 1-9 and 1 ImageNet session for subjects 10-30), where each image represented one of the 1,000 ImageNet categories with no overlap across categories. The authors did not define a training and testing split. Here, we defined a 80%/20% training and testing split per subject by assigning the top 20% most perceptually diverse ImageNet and the top 20% most perceptually diverse COCO images to the test set. The perceptual diversity of image X was measured as the median of the distances (1-cosine similarity) between image X’s DreamSim embedding [42] and every other image’s DreamSim embedding presented to that subject. Since all NOD subjects viewed unique ImageNet images, each subject has a different set of training and testing set images. The training and testing set split for the 120 COCO images were shared between the first nine NOD subjects that viewed them.

#### 7.2.7 Natural Scenes Dataset

The Natural Scenes Dataset (NSD) [11] was collected in Minneapolis, MN USA. Subjects 1-8 participated in 40, 40, 32, 30, 40, 32, 40, and 30 core experimental sessions, respectively. Naturalistic images in the core experimental sessions were presented with 3 second duration at 8.4 degrees of visual angle overlaid with a center fixation dot. A 1 second fixation period followed the offset of the image presentation. Subjects were instructed to press a button with their right index finger if the observed image was new or press a second button with their right middle finger if the observed image was old. Button presses were recorded between 250 and 4,250 milliseconds post-stimulus onset. Five out of 68 trials per run were blank trials.

Subjects also participated in one synthetic session where they viewed synthetic stimuli (such as colors, textures, line drawings, words, upside down images) with a 2 second duration followed by a 2 second blank screen (4 second trial) [133]. The vigilance task switched between fixation and one-back in alternating runs.

NSD’s stimulus set contains 70,566 unique naturalistic images from the Common Objects in Context (COCO) database [125] and 284 artificial images [133]. A naturalistic testing set of 1,000 images were shared across subjects and presented one to three times per subject. A naturalistic training set of 69,566 images were also shown one to three times per subject but were not shared between subjects. In total, a subject viewed between 9,000 and 10,000 unique naturalistic images over the course of the core experiment. The artificial testing set of 284 images were presented 2-8 times each. Each subject viewed 64 unique color textures calibrated to their eyesight, but given the stimuli’s high similarity across subjects, we averaged the RGB values over subjects and treated these 64 stimuli (along with the remaining 220 synthetic stimuli) as ’shared’ across subjects.

Each NSD subject also underwent between 100 and 180 minutes of resting-state fMRI for a total of 960 minutes (16 hours) of resting-state data in the dataset. Subjects were instructed to fixate on a white fixation cross on a gray background, rest, and stay awake. Some resting-state runs included additional instructions to inhale deeply upon changing color of the fixation cross to aid in physiological data analysis.

#### 7.2.8 THINGS fMRI

THINGS-fmri (THINGS) [48] was collected in Bethesda, Maryland USA. The 3 subjects participated in 12 main experimental sessions. Each image trial consisted of an image presentation of 0.5 second duration at 10 degrees of visual angle overlaid with a center fixation cross followed by a 2.5 second fixation period. Subjects were instructed to press a button when they viewed a synthetic image. The 100 test images were presented exactly once in random order throughout each session.

The THINGS dataset sampled 8,740 naturalistic images belonging to 1 of 720 nameable object concepts. 8,640 training set images were presented once per subject and 100 testing set images were presented 12 times per subject. The images were derived from the ImageNet database [127], Google images, and Bing images.

Subjects 1, 2, and 3 underwent 84, 84, and 78 minutes of eyes-closed resting-state fMRI scans, respectively.

### 7.3 Stimuli aggregation

Stimuli were aggregated across all 8 datasets and filtered to account for (1) testing and training set conflicts across datasets, and (2) perceptually similar stimuli between train and test splits (expanded later). The final dataset contains 162,839 unique stimuli partitioned into a naturalistic training set of 144,360 stimuli (353,924 single trial responses), a naturalistic testing set of 18,145 stimuli (68,011 single trial responses), and an artificial testing set of 334 stimuli (8,072 single trial responses). We summarize each dataset’s stimulus set in Table 2 and in the text below.

In this study we process and provide all stimuli and their associated brain responses described above but define training and testing subsets of the stimuli for analyses that depend on the split. When compiling the stimuli across datasets, we prioritize preserving each dataset’s original train-test splits and retaining the highest number of brain responses while reducing the perceptual similarity between the test and train splits.

We first naively group all train and test stimuli according to their dataset-specific training and testing splits described above. No distinction between naturalistic and artificial stimuli were made at this stage. We then impose two filters to create a perceptually distinct final training and testing set. The first filter resolves conflicts where the same stimulus by filename is in both the training set of one (or more) dataset(s) and in the testing set of one (or more) datasets. This case affects 1,032 stimuli. The stimulus and associated trials belonging to the set with the least number of total single trials was removed. In the case of a tie, the stimuli and trials belonging to the test set is preserved. The second filter ensures that no training set stimulus is too similar to a testing set stimulus by discarding training set stimuli and their associated trials that exceed a predefined similarity threshold (cosine similarity between DreamSim model embeddings [42]) to a testing set stimulus. A threshold of 0.8196 was determined by computing the average similarity between the first and last video frames over all 1,102 video stimuli from the BOLD Moments Dataset. Since the BOLD Moments Dataset stimuli contain no cut scenes, the similarity between the first and last frames provide a good similarity heuristic of the same visual content that differs primarily in camera angle and object placement. This similarity threshold effectively excludes training stimuli that can reasonably be perceived as being from the same scene or of the same object as a testing set stimulus. This second filter discards a total of 1,186 training stimuli that were too similar to one of 622 testing stimuli (i.e., some testing stimuli contained more than one highly similar training stimuli). In the case of BMD and HAD videos, embeddings were averaged over frames.

The training and testing splits are defined by the stimuli, not brain responses. Rather, from a modeling point of view, brain responses are treated as a label to a stimulus. Thus, a single stimulus filename in either the training or testing split can be associated with multiple single trial brain responses across subjects and datasets.

This process preserves a stimulus’s original train-test set assignment to allow fair comparisons with prior work using each dataset author’s proposed train-test splits, when available. While this study’s train-test split may exclude some stimuli compared to the original split, the stimuli are not mixed between splits. Crucially, the filters control for test-train leakage by removing highly similar or even duplicate stimuli (regardless of the stimulus filename) and thus enable strong tests of generalizability.

### 7.4 fMRI dataset preprocessing

#### 7.4.1 Task-fMRI preprocessing

The fMRI datasets were downloaded from their original source (see Data and Code Availability statement) after their respective dicom to Nifti conversion performed by the dataset authors. We then use fMRIprep (version 23.2.0) to preprocess and register the data to fsLR32k cortical surface space, preferred for its superior inter-subject registration over volumetric registrations [52, 59, 134] and seamless integration with Human Connectome Project tools [135] and the HCP-MMP1.0 parcellation [43] (see Region of Interest section below). All fMRI data were slice-time corrected to the first slice in the acquisition period and the functional bold scans were registered to the T1w structural scan with 12 degrees of freedom. We then estimate single trial beta values for each subject separately using a General Linear Model (GLM) [41].

GLMsingle first fits a Hemodynamic Response Function (HRF) to each voxel (GLMsingle types B, C, and D). When multiple repetitions of a stimulus exist, it then employs a cross-validation procedure to estimate noise and optimally chooses up to 10 PCA components as nuisance regressors (GLMsingle types C and D). Finally, it uses fractional ridge regression to achieve more accurate beta estimates, especially in event-related designs used here where the BOLD signal may overlap between trials (GLMsingle type D).

For each dataset where the acquisition TR was not time-locked with stimulus onsets and duration (BMD, BOLD5000, NSD, NOD, THINGS), the time series at each vertex was resampled (linearly interpolated) to the nearest TR that is a multiple of the stimulus onset and offset [136]. For example, the NSD’s acquisition TR of 1.6 seconds was temporally resampled to a TR of 1 second to accommodate its trial length of 4 seconds and stimulus duration of 3 seconds. Next, the GLM was computed for each session independently (BMD, NSD, HAD, NOD subjects 10-30) or, when no stimuli were repeated within a session but were repeated across sessions, for multiple sessions concatenated together (BOLD5000, THINGS, GOD, Deeprecon, NOD subjects 1-9). The session concatenation was performed when possible to take advantage of GLMsingle’s type D procedure because its resulting betas estimates show significantly higher noise ceilings than type B’s beta estimates [41]. When multiple sessions were concatenated together, GLMsingle’s ’session indicator’ option was used to internally z-score the values over each vertex to account for differences in magnitude between sessions. In the case of HAD subjects 1-30 and NOD subjects 10-30, where each stimulus was repeated exactly once throughout the experiment, six motion regressors (X, Y, and Z translation and rotation) were used as nuisance regressors since cross-validation could not be performed to determine PCA nuisance regressors.

The beta estimates output by GLMsingle were returned in units of percent signal change. They were then normalized within each session (or group of sessions) by first identifying the training set stimuli that were presented within that session (or group of sessions). The mean and standard deviation at each vertex for those training set stimuli was computed. Then, normalized beta estimates at each vertex were computed by subtracting the training set mean and dividing by the training set standard deviation from both the training and testing sets. This normalization procedure (as opposed to z-scoring training and testing sets independently) better accommodated datasets where each session (or group of sessions) had few testing set examples. The training set stimuli used in the normalization were those defined with respect to the individual dataset (detailed above), not the dataset compilation. No spatial smoothing was applied to the data.

Trials were only discarded from this present study if their corresponding stimulus was discarded in the stimulus set filtering process (see Table 2). Below we describe dataset-specific GLM estimation details.

##### BOLD5000

Due to a lack of within-session stimulus repetitions, we concatenated sessions 1-5, 6-10, and 11-15 for subjects 1-3 and sessions 1-5 and 6-9 for subject 4 when estimating single trial beta values [41]. In this way, we take advantage of GLMsingle’s type D beta estimates. All functional runs were temporally interpolated from the acquisition TR of 2 seconds to a TR of 1 second. The stimulus duration was set to 1 second. The resulting beta estimates were then normalized by the mean and standard deviation of the original training stimuli’s values within that session grouping.

##### BOLD Moments Dataset

For each subject, the GLM was computed for each of the 4 core experimental sessions separately. The fMRI time-series was linearly interpolated from the acquisition TR of 1.75 seconds to a TR of 1 second. Stimuli were modeled with a duration of 3 seconds. Blank trials were not included in the GLM. GLMsingle type D procedure was used to estimate the single trial beta values. The beta estimates were then normalized by the mean and standard deviation of the dataset’s pre-specified training split within each session.

##### Human Actions Dataset

For each subject, we compute single trial beta estimates over the subject’s 1 session. Since there were no stimulus repeats within each subject, we use GLMsingle’s type B trial estimates and input six rigid body motion confounds (x, y, z rotation and translation) as regressors of no interest. Since the vigilance task was highly correlated to stimulus features (pressing a button with the right and left thumb if the video depicted a ’sport’ or ’non-sport’ action, respectively), two additional nuisance regressors were input to the GLM corresponding to the left and right thumb button presses. Button press regressors were computed by convolving a canonical HRF (SPM) with the button press onsets, modeled with 0 second duration, within the run. The stimulus duration was modeled as 2 seconds, the length of the video. The resulting single trial beta estimates were normalized by the training set’s (stimuli in the first 9 runs, described above) mean and standard deviation.

##### THINGS fMRI

Since no stimuli were repeated within a session, we concatenate sessions 1-6 and 6-12 to estimate single trial betas using GLMsingle’s type D procedure for each subject separately. The fMRI time-series was temporally interpolated from the acquisition TR of 1.5 seconds to 0.5 seconds. The stimulus duration was modeled as 0.5 seconds. The synthetic image catch trials were not modeled in the GLM. The beta estimates were then normalized within each session group by the training set stimuli’s mean and standard deviation.

##### Natural Scenes Dataset

A GLM (type D) was computed for each session independently. The fMRI time series input into the GLM was temporally resampled from the acquisition TR of 1.6 seconds to a TR of 1 second to evenly fit into the stimulus duration of 3 seconds and total trial length of 4 seconds. The beta estimates for each session were then normalized by the mean and standard deviation of the training stimuli occurring within that session.

##### Natural Object Dataset

Since subjects 1-9 viewed multiple repetitions of COCO images during the COCO session, the 4 ImageNet sessions and the COCO session were concatenated together to estimate single trial beta estimates using GLMsingle’s type D procedure. Trials used for the one-back vigilance task in the COCO sessions were not input into the GLM. Subjects 10-30 viewed one ImageNet session with no repeated stimuli and thus used GLMsingle’s type B procedure to estimate single trial betas. For the type B procedure, six rigid body motion confounds (x, y, and z rotation and translation) were input into the GLM as regressors of no interest. Due to the high correlation between the vigilance task and stimulus features (pressing a button with the right and left thumb if the image depicted ’animate’ or ’inanimate’ content, respectively), we add two additional nuissance regressors to the GLM to model the left and right thumb button presses. Button press regressors for each run (ImageNet runs only) were modeled with button press onsets at 0 second duration and convolved with a canonical HRF (SPM). The fMRI time series for all subjects was temporally interpolated from the acquisition TR of 2 seconds to 0.5 seconds. The stimulus duration was modeled as 0.5 seconds for both the ImageNet and COCO sessions despite the ImageNet images being presented for 1 second. Specifying the same stimulus duration across the ImageNet and COCO sessions was necessary in order to use GLMsingle’s type D beta estimation procedure. For consistency, we also specified a 0.5 stimulus duration for the ImageNet session in subjects 10-30. The resulting beta estimates were normalized according the mean and standard deviation of the training and testing set stimuli.

##### Generic Object Decoding

All three naturalistic training (no stimulus repeats) and 4 naturalistic testing sessions (contained stimulus repeats) were concatenated together as input for GLMsingle’s type D beta estimation. The duration of the stimulus was specified as 9 seconds. The resulting beta estimates were normalized by the training set’s mean and standard deviation.

##### Deeprecon

The naturalistic training, naturalistic testing, artificial shape, and artificial letter stimuli were all presented within stimulus-specific sessions and repeated within subjects. Thus, the sessions corresponding to each of the 4 groups were concatenated together and single trial beta values were estimated with GLMsingle’s type D procedure. The stimulus duration was specified as 8 seconds for the naturalistic training, naturalistic testing, and artificial shape session groupings. The stimulus duration was specified as 12 seconds for the artificial letter session grouping. One-back vigilance task trials and resting blocks (for the artificial letter sessions) were not input into the GLM. The beta estimates within each session grouping were normalized by the mean and standard deviation computed across all beta estimates since the session groupings did not mix train and test splits.

#### 7.4.2 Resting state and task fMRI time-series preprocessing

After preprocessing all fMRI data with fMRIPrep, the full runs of resting state (applicable to BMD, NSD, THINGS) and task fMRI data were independently cleaned using Nilearn’s clean img function. Following recommendations applied to the Human Connectome Project resting state data [59, 60], we detrend, filter (high pass = 0.008 Hz, no low pass filter applied), regress out confounds, and standardize to unit variance (in that order) of the data. For the confound regression, we use a set of 24 timeseries confounds derived from fMRIPrep’s motion correction procedure: X, Y, and Z translation and rotation, their temporal derivatives, plus the square of those 12 regressors. The Connectome Workbench software package was used to convert the fsLR32k cortical surface to volume space to apply the cleaning and then to convert the cleaned image back to fsLR32k surface space. This time-series data is independent of the beta-stimulus pairs used in the above analyses. We release these time-series data to encourage research directions that integrate the Human Connectome Project’s resting state fMRI [59, 137] (the s1200 release adds an additional 60 minutes of resting state scans for 1,018 subjects) or explore task-based encoding/decoding approaches on time-series data [13].

### 7.5 Vertex-wise noise ceiling estimation

For datasets and subjects with multiple stimulus repetitions (all datasets and subjects minus HAD and NOD subjects 10-30), we estimate each subject’s noise ceiling at each vertex using the single-trial beta values as brain responses [11]. We assume that repeated viewings of the same stimulus should evoke the same fMRI response, and a response to a stimulus is a combination of stimulus-induced signal and Gaussian noise (regardless of whether its source be scanner, neural, physiological or other) [138]. Subject-specific noise ceilings at each vertex were estimated for each evaluation set: naturalistic train, naturalistic test, and artificial test, where applicable. Within each subject and evaluation set, a single-trial and trial-averaged noise ceiling was computed. To summarize the procedure, we first derived the signal variance by subtracting the data’s total noise variance from its total variance. We then computed a noise ceiling SNR (*ncsnr*) term as the ratio of signal to noise standard deviations. The noise ceiling, in terms of variance explained, is *ncsnr*^2^ divided by *ncsnr*^2^ + *t_avg_*, where the *t_avg_* term modulates the noise ceiling estimation by how many trials are being averaged. We assume that the beta response data for a given subject and evaluation set may contain stimuli with varying number of repetitions.

To calculate the total noise variance, we pool the noise variance across subsets of the data with equal number of repetitions per stimulus by weighting each subset’s noise variance by its degrees of freedom.

For a given data subset *p*, the noise variance was computed as:

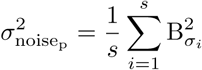

where *s* is the number of stimuli in the subset and 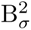 is the variance between stimulus repetitions using *n* − 1 degrees of freedom.

Pooling each subset’s noise variance gives the total noise variance of the data 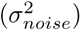

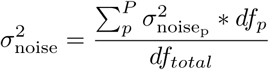

where *B* is the number of data subsets, *df_p_* is the degrees of freedom for the *p*’th subset (*nstimuli* ∗ (*nrepetitions* − 1)), and *df_total_* is the sum of each subset’s degrees of freedom. Note that if all stimuli across all the data have the same number of repetitions, there are no subsets and the degrees of freedom terms cancel.

We then compute the total variance for each vertex as square of the standard deviation over all single trial beta estimates from all stimuli and repetitions:

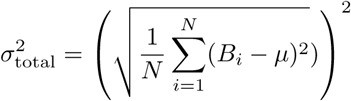

where *B* is a single trial beta estimate, *µ* is the average response at the vertex, and *N* is the number of single trials.

The signal variance is estimated as total variance minus the total noise variance:

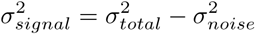

The signal variance was positively rectified where negative values were mapped to 0 and positive values remained the same. The rectified signal variance and the noise standard deviation was used to compute a noise ceiling SNR measure:

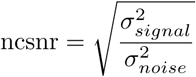

Finally, we use *ncsnr* to compute a noise ceiling measure expressed as percent of signal variance to total variance for different degrees of trial averaging:

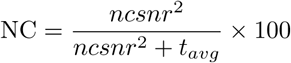

To estimate a noise ceiling for a constant number *n* of averaged trials per stimulus, the *t_avg_* term is 1*/n*. In the case where stimuli have varying number of repeats:

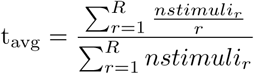

where *nstimuli_r_*is the number of stimuli with *r* repeats and *R* is the maximum number of repetitions of a stimulus in the data.

### 7.6 Regions of interest

We use the HCP-MMP1.0 parcellation [43] to define Regions of Interest (ROI). The HCP-MMP1.0 parcellation is defined by functional, connective, topographic, and architectural properties across 210 healthy adult subjects from the Human Connectome Project [9]. The parcellation defines a total of 180 cortical regions per hemisphere. Geographically contiguous regions that share functional, connective, topographic, or architectural properties are additionally assigned to one of 22 super groups. The HCP- MMP1.0 parcellation’s ROI definitions are accurately registered to the fsLR32k space as both the parcellation and the fsLR32k space are derivatives of the Human Connectome Project. We access the ROI definitions for analysis and visualizations using the hcp-utils (GitHub: https://github.com/rmldj/hcp-utils) Python package. Since all functional fMRI data in this study is also registered to the fsLR32k space, the HCP-MMP1.0 parcellation enables accurate vertex-to-vertex and ROI-to-ROI correspondences across subjects and datasets. Note that we do not use subject-specific ROI definitions (e.g., those defined via functional localizers or retinotopic experiments) in this study.

The fsLR32k space [52] uses surface-based registration to define cortex and a volume-based registration to define subcortical structures. The fsLR32k mesh is composed of 91,282 ”grayordinate” points, where 29,706 vertices define the left hemisphere cortex, 29,706 vertices define the right hemisphere cortex, and 31,870 voxels define subcortical structures. Cortical vertices are sampled roughly 2mm apart and subcortical voxels are defined with 2mm voxel resolution. In this work, only the left and right hemisphere cortical vertices, fully specified by the HCP-MMP1.0 parcellation’s 180 per-hemisphere ROIs, are analyzed.

Various sources of noise, such as fMRI scanner dropout, subject motion artifacts, and fMRI scanner inhomogeneities, may lead to undefined vertices (e.g., values of NaN or Inf) in individual fMRI trials. While removing vertices from all subjects that were undefined in potentially only one trial of one subject guarantees that the undefined vertices will not confound results, we recognize that some analyses may benefit from using the most signal when available or require all vertices to take on a finite value. To accommodate such analyses, we control for the undefined vertices for each single trial brain response by filling in their value with the mean value of immediately adjacent cortices. Specifically, for each undefined vertex, we calculate its fraction of adjacent vertices that are defined by a finite value. We identify the vertex with the greatest fraction of defined neighbors and fill in its value as the mean of its defined neighbors. We then recalculate the fraction of well-defined adjacent vertices for each remaining undefined vertex and repeat until no more undefined vertices remain. This algorithm does not discard any vertices and preserves the well-known spatial correlation between cortical responses, as opposed to other methods that might fill in the undefined vertices with single values (i.e., all zeros) or a value sampled from a distribution. The goal of this procedure is not to estimate true signal at undefined vertices but rather to preserve signal at vertices that are undefined in some trials while aiming to minimize ’unique identifiers’ in the data from stark differences in distributions. Ultimately, any technique to fill in missing values may unintentionally introduce biases, so ensuring results are interpretable and generalizable are paramount.

Across all trials from all subjects and datasets, a total of 2,361 cortical vertices (out of 59,412 cortical vertices) were undefined in at least one trial. We show which subjects and ROIs had trials with undefined vertices in Supplementary Fig. A2.

### 7.7 Decoding from high and low diversity stimulus subsets

High and low diversity stimulus subsets were selected by first randomly choosing one of the stimuli observed by the subject as a seed. All pairwise distances were then computed between the seed’s DreamSim [42] features and every other stimulus’s DreamSim features. The 200 nearest neighbor stimuli were selected as the low diversity subset. The high diversity subset was selected using the farthest-first algorithm, where, starting from the stimulus furthest away from the seed, the stimulus with the maximum minimum pairwise distance to the already selected high diverse stimuli is added to the subset until 200 stimuli are sampled. The same stimulus can be part of both the high and low diversity subsets. Given the high and low diversity subsets become increasingly similar as the number of samples increases, we fixed the number of samples to 200 to balance maintaining a strong distinction between subsets and sampling a reasonable amount of stimuli.

A linear decoding model (ridge, k=5 fold cross validation, best *α* selected log-linearly between 10*^−^*^4^ and 10^8^) was trained to map brain activity from the visual cortex (MMP1.0 sections 1-5) to the corresponding DreamSim features. PCA (n=100 features) and normalization (0 mean, unit standard deviation) was fit to the training set brain activity and applied to the testing set brain activity before model training. Identical PCA and normalization was also applied to the DreamSim features before model training. All models were trained on data from one subject at a time. One hundred PCA components was chosen to ensure that the number of features was less than the number of samples (n=200).

The models trained on the high diversity and low diversity subsets each predicted the DreamSim features of each subject’s naturalistic test set. Prediction accuracy was computed by correlating (Pearson) the predictions with the ground truth DreamSim features (after PCA) and averaging over the samples in the test set. The entire procedure from stimulus subset sampling to model prediction was repeated fifty times per subject in order to obtain a distribution of predictions and control for undesired confounds between a subset’s stimulus features and target features.

Decoding was also performed with varying levels of training data (Fig. 4f). PCA was applied to the brain activity and DreamSim features in the same way described above. The number of samples was ensured to be at least 200 to allow a reasonable ratio between number of samples and PCA features. The farthest-first sampling strategy and the nearest neighbor sampling strategy are described above for the high and low diversity sets, respectively, but here the exact number of stimuli in the final set is varied. The random sampling method randomly chose the desired number of samples without replacement. Decoding results were also repeated fifty times per training set size and sampling strategy to measure the variation in the data. To ensure the decoding results from the three sampling strategies were equal when using 100% of the data, the indices were sorted so the cross-validation procedure would select the same hyperparameters.

### 7.8 Brain-optimized model architectures

The CNN8 model architecture followed that used in [12] and was composed of a shared core network consisting of eight 2D convolutional blocks followed by subject-specific linear factorized readout heads [95]. Each convolutional block consisted of 2D convolution, 2D batch normalization, and ReLU layers. The convolutional layer kernel sizes ranged from 5x5 (blocks 1 and 2) to 3x3 (blocks 3-8) and padding ranged from 1 (blocks 1, 3-8) to 2 (block 2). No bias was used in the convolutional layers. The number of input and output channels for each convolutional layer was 384 (except the 3 RGB channel features for the initial input). Convolutional weights were initialized with kaiming uniform initialization. An average pooling layer of kernel size 2 and stride 2 was inserted after blocks 2, 4, and 6. An average pooling layer of kernel size 2 and stride 1 was inserted after block 8. Each readout head consisted of two weight matrices (”spatial” and ”features” matrices) initialized with a bias of 0 and weights following a normal distribution (mean 0, std 0.001). The spatial weights were normalized with L2 normalization.

We additionally used the PyTorch implementation of AlexNet [97], SqueezeNet1.1 [98], ResNet18 [99], and Swin-T [100] models (Fig. 5d). To adapt these models from image classification to brain activity prediction, the final layer(s) that flattened the feature map for the purpose of image classification were discarded so the output (i.e., input to the linear readout head) was shape (channels x height x width). AlexNet and SqueezeNet1.1 architectures additionally added BatchNorm2D layers after each convolutional layer to stabilize brain-optimized training and improve brain prediction performance. The task-optimized versions of each of the four models used the publicly available IMAGENET1k V1 model weights from PyTorch.

### 7.9 Brain-optimized model training

Models were trained on seven Nvidia Titan RTX GPUs for a maximum of 100 epochs. Early stopping was used if no improvement to validation loss (above delta=0.002) was recorded for 7 epochs. Model weights resulting in the lowest validation loss at the end of an epoch were saved. The learning rate was initialized a 1e-4 and was scheduled to decrease by a factor of 0.3 if no relative improvement (threshold=1e-4) in validation loss was seen after 6 epochs (PyTorch’s ”ReduceLROnPlateuau”). The model was trained with an Adam optimizer initialized with a weight decay of 1e-4. The model weights were initialized with the default kaiming initialization. The model was trained with a batch size of 64 composed of a random sampling of fMRI-stimulus pairs across all 93 subjects (unless stated otherwise). The loss function was mean squared error (MSE). The train/validation split was 90%/10%, respectively.

Stimulus inputs were center square cropped along the shorter dimension, resized to 224x224 pixels, and normalized (mean=[0.485, 0.456, 0.406], std=[0.229, 0.224, 0.225]). For video inputs (i.e., BMD and HAD), the middle frame was used. No data augmentations were used as standard data augmentations (e.g., flipping, rotating) could reasonably result in a different brain response label and thus add noise to the training.

The fMRI data per stimulus was averaged over all trials for a given subject to increase signal. The model predicted a final output of 7,831 vertices (unless stated otherwise) corresponding to defined values in the MMP1.0 parcellation sections 1-5 [43]. No other vertex selection criteria, such as a noise ceiling threshold, was used.

### 7.10 In silico functional localizer

We perform the localizer experiment described in [106] on a brain-optimized model composed of a 8-layer CNN (CNN8) core and eight subject specific readout heads for the eight Natural Scenes Dataset subjects. The model was not trained on all 93 MOSAIC subjects due to the each readout head’s larger size from whole brain prediction (57,051 non-nan vertices) that exceeded our available GPU memory. Once trained, we run inference on the localizer’s set of 1,584 images for each subject separately. The images underwent the same image transforms used in training, where they were center square cropped along the shorter dimension, resized to 224x224 pixels, and normalized (mean=[0.485, 0.456, 0.406], std=[0.229, 0.224, 0.225]).

Six contrasts were performed between the different image categories following the protocol outlined in [106] to identify character-, body-, face-, place-, object-, and object LOC-selective regions (character: [’word’, ’number’] *>* [”body”, ”limb”, ”child”,”adult”, ”corridor”, ”house”, ”car”, ”instrument”], body: [”body”, ”limb”] *>* [”child”,”adult” ,”corridor”, ”house”, ”car”, ”instrument”, ”word”, ”number”], face: [”child”, ”adult”] *>* [”word”, ”number”, ”body”, ”limb”, ”corridor”, ”house”, ”car”, ”instrument”], place: [”corridor”, ”house”] *>* [”word”, ”number”, ”body”, ”limb”, ”child”, ”adult”, ”car”, ”instrument”], object: [”car”, ”instrument”] *>* [”word”, ”num-ber”, ”body”, ”limb”, ”child”, ”adult”,”corridor”, ”house”], object LOC: [”car”, ”instrument”] *>* [”scrambled”]). For each contrast, the average responses of the negative categories were subtracted from the average responses of the positive categories. A contrast threshold was determined by computing the maximum F1 score from a precision-recall curve. All vertex values above the threshold defined the predicted binary contrast. The selected set of vertices were not constrained in any way, such as to be contiguous or within a certain cortical region or hemisphere.

We then used the trial-averaged fMRI responses from a given subject and their category-specific masks to identify the top 100 stimuli with the highest vertex-averaged response within the category-specific mask. Both naturalistic and artificial stimuli were able to be chosen.

### 7.11 Statistics

In section 2.2, we use the non-parametric Wilcoxon signed-rank test [139] to test significant differences between high minus low diversity subset correlations (Fig. 4d) and high minus low diversity subset standard deviation (Fig. 4e) (p*<*0.05, two-sided, against a difference of zero). Effect sizes were estimated using the Common Language Effect Size (CLES) expressed as *n positive* + (*n ties/*2))*/n total*.

To measure the subject-specificity of the fROIs, we also use the Common Language Effect Size expressed as *n positive* + (*n ties/*2))*/n total*. In other words, for each subject, it is the proportion out of four contrasts where the same-subject is higher ranked than the mean of the other-subject ranks.

## Appendix A Appendix

**Table A1:**
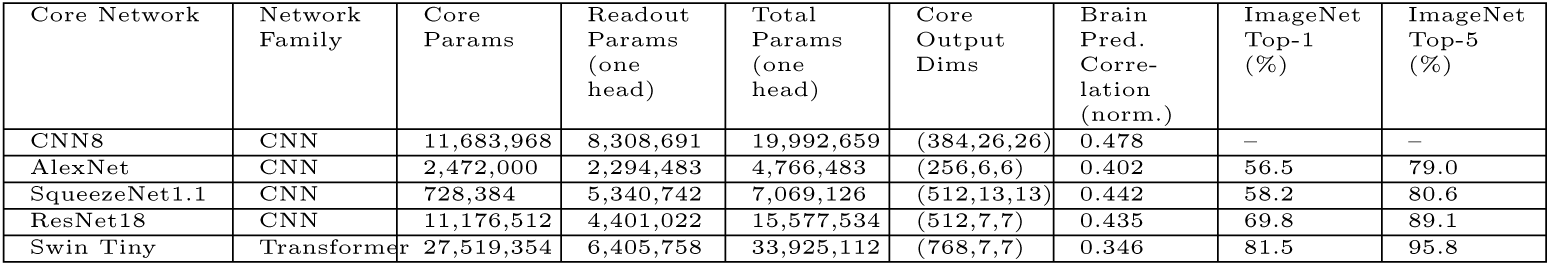
Differences in brain-optimized architectures. Brain-optimized networks are constructed using different core network architectures to predict visual cortex (MMP1.0 sections 1-5, 7831 output vertices). The Brain Prediction Correlation (normalized) column averages NSD subjects 01-08 and Deeprecon subjects 01-03 median voxelwise correlation over visual cortex vertices predicted with the 93-subject multi-head framework. The ImageNet1k task accuracies use the respective model’s pretrained weights available through PyTorch. Note that the number of parameters in the brain-optimized core network may differ from its ImageNet1k-optimized counterpart due to the trimming of fully connected layers and, in the case of AlexNet and SqueezeNet1.1, the addition of BatchNorm2d layers.

| Core Network | Network Family | Core Params | Readout Params (one head) | Total Params (one head) | Core Output Dims | Brain Pred. Correlation (norm.) | ImageNet Top-1 (%) | ImageNet Top-5 (%) |
| --- | --- | --- | --- | --- | --- | --- | --- | --- |
| CNN8 | CNN | 11,683,968 | 8,308,691 | 19,992,659 | (384,26,26) | 0.478 | – | – |
| AlexNet | CNN | 2,472,000 | 2,294,483 | 4,766,483 | (256,6,6) | 0.402 | 56.5 | 79.0 |
| SqueezeNet1.1 | CNN | 728,384 | 5,340,742 | 7,069,126 | (512,13,13) | 0.442 | 58.2 | 80.6 |
| ResNet18 | CNN | 11,176,512 | 4,401,022 | 15,577,534 | (512,7,7) | 0.435 | 69.8 | 89.1 |
| Swin Tiny | Transformer | 27,519,354 | 6,405,758 | 33,925,112 | (768,7,7) | 0.346 | 81.5 | 95.8 |

**Fig. A1:**
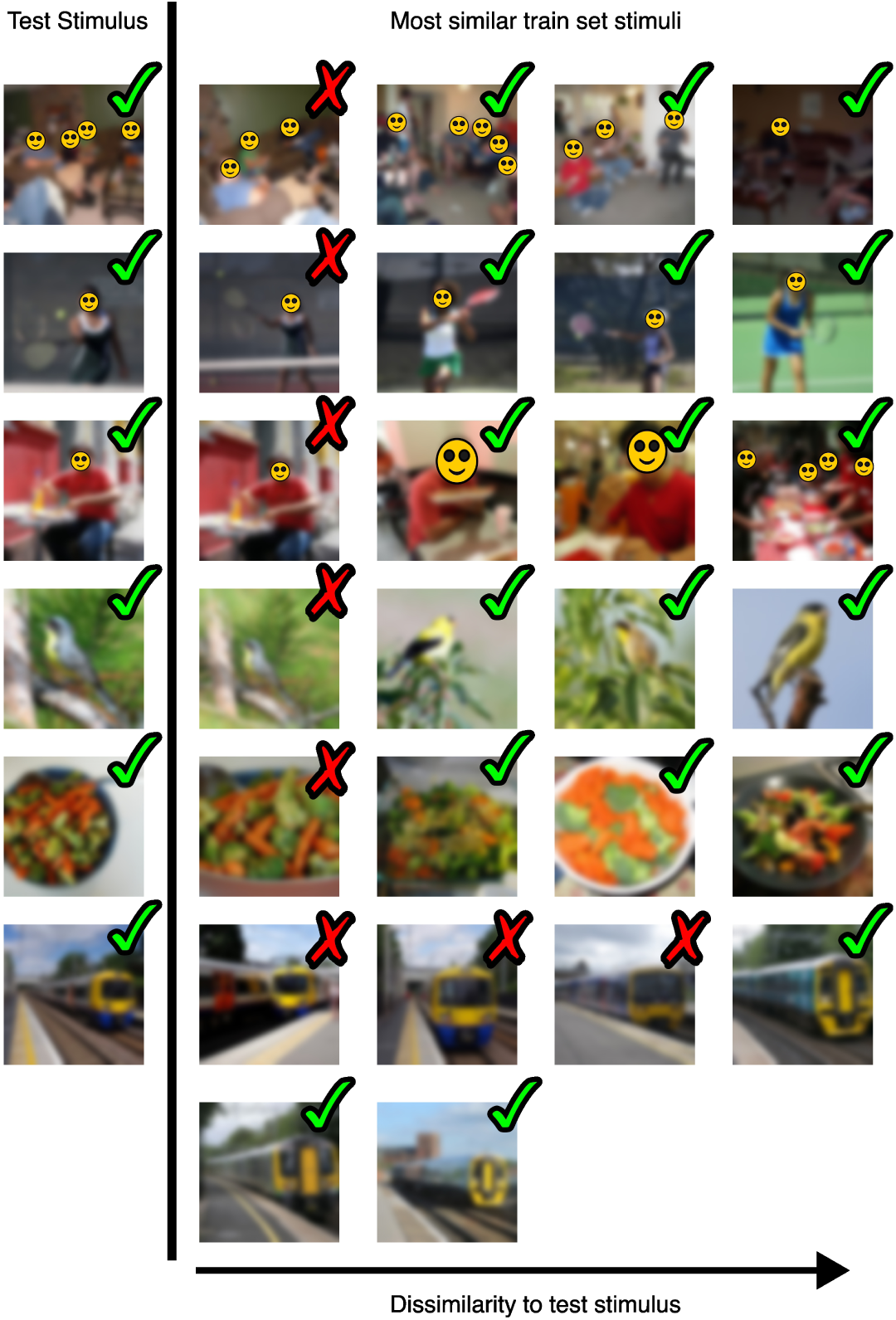
Examples of stimulus curation procedure. The perceptual similarity, as measured by cosine similarity between each stimulus’s DreamSim embeddings, was computed between each test stimulus and all train stimuli. Train stimuli that exceed a predetermined similarity threshold are excluded from the train set as to limit train-test set leakage in the aggregated MOSAIC dataset. The graphic shows the most similar train stimuli to a test stimulus and whether the train stimulus exceeded the threshold to be excluded (red X) or not (green checkmark). Note that stimuli that fall below the similarity threshold for a given test set stimulus (green checkmark) can still be excluded for being too similar to another test stimulus. The first two rows depict excluded train stimuli that are a different moment in time of a test stimulus. The middle two rows show excluded train stimuli that are crops of a test stimulus. The final two rows show excluded train stimuli that are different scenes from a test stimulus but are perceptually very similar. Images are blurred and identifiable faces are masked for copyright and privacy purposes. The unedited figure is available with the data release.

**Fig. A2:**
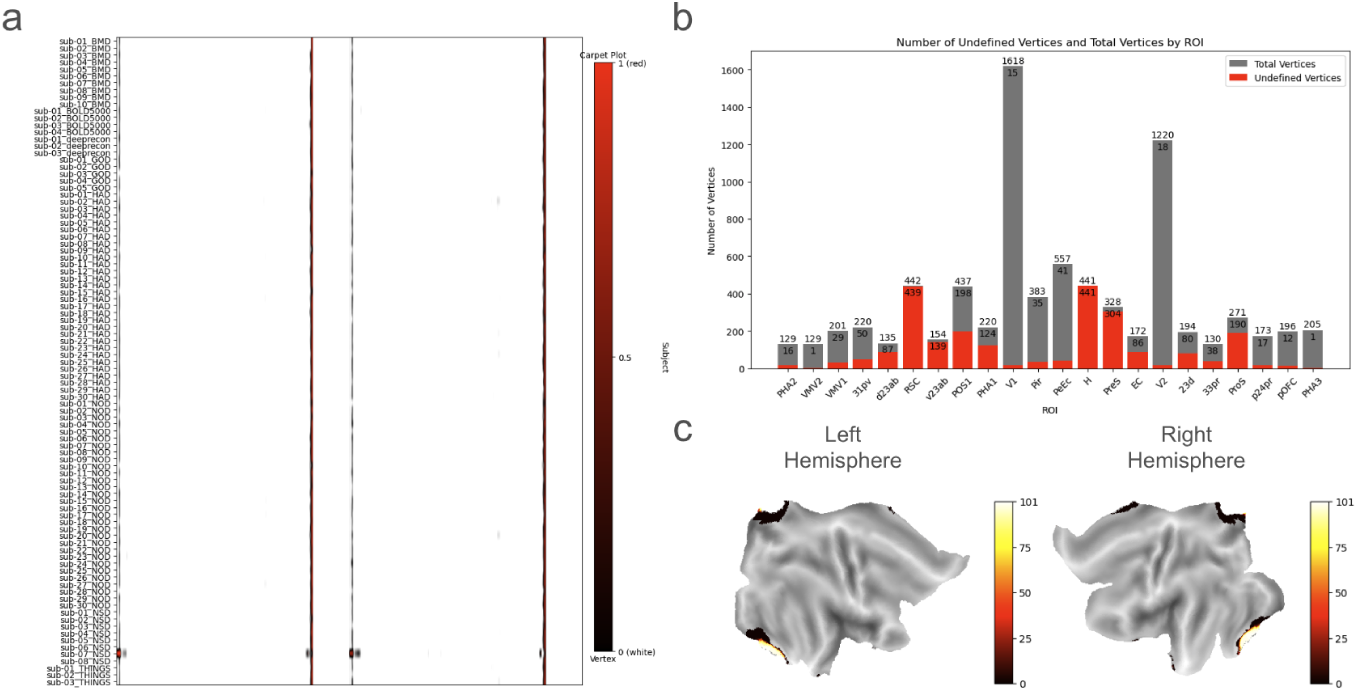
Undefined cortical vertices across subjects. **a** Undefined vertices across subjects: The carpet plot highlights the percent of trials with undefined values at each vertex for each subject. The color white signifies the vertex for that subject was never undefined. If the vertex were undefined at at least one trial, the percent of trials with undefined values is colored according to the black-red gradient. **b** Undefined vertices per ROI: For each region of interest (ROI) with at least one vertex undefined over all trials and subjects, the gray bar shows the total number of vertices (exact number above the bar) and the red bar shows the number of undefined vertices (exact number below the bar). **c** Location of undefined vertices: The average of subject-average undefined values divided by number of trials is plotted on the flat map. A color of gray denotes the vertex was never undefined over all trials and subjects. The black to white gradient shows how often the vertex was undefined.

**Fig. A3:**
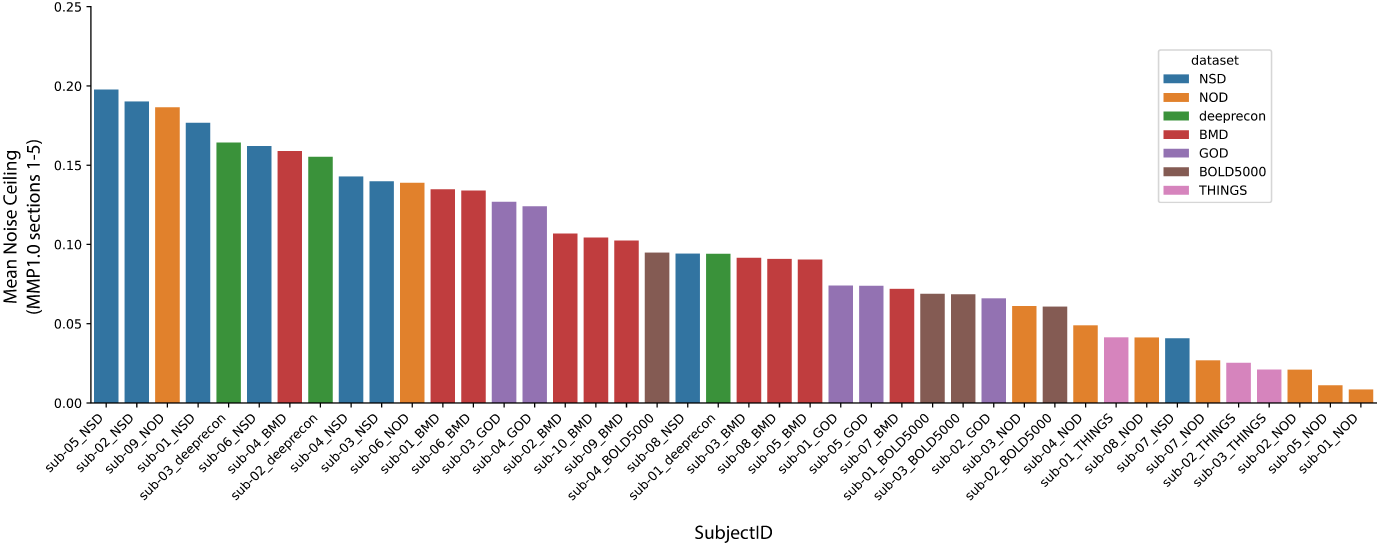
Average noise ceiling in visual cortex by subject. The noise ceiling at each vertex was estimated for the 43 subjects that contained repeated stimulus presentations (all subjects except Natural Object Dataset subjects 10-30 and Human Actions Dataset subjects 1-30) as described in 7.5. Here we plot the noise ceiling averaged over visual cortex (MMP1.0 sections 1-5, 7831 vertices) for each subject. Single-trial noise ceiling estimates (n=1) were computed from responses in each subject’s naturalistic train set, when available. Because subjects in Generic Object Decoding, THINGS, and BOLD5000 datasets were not shown repeated stimuli in their naturalistic train set, responses in their naturalistic test set was used. Note that these average noise ceiling estimates are only a coarse approximation of data quality.

**Fig. A4:**
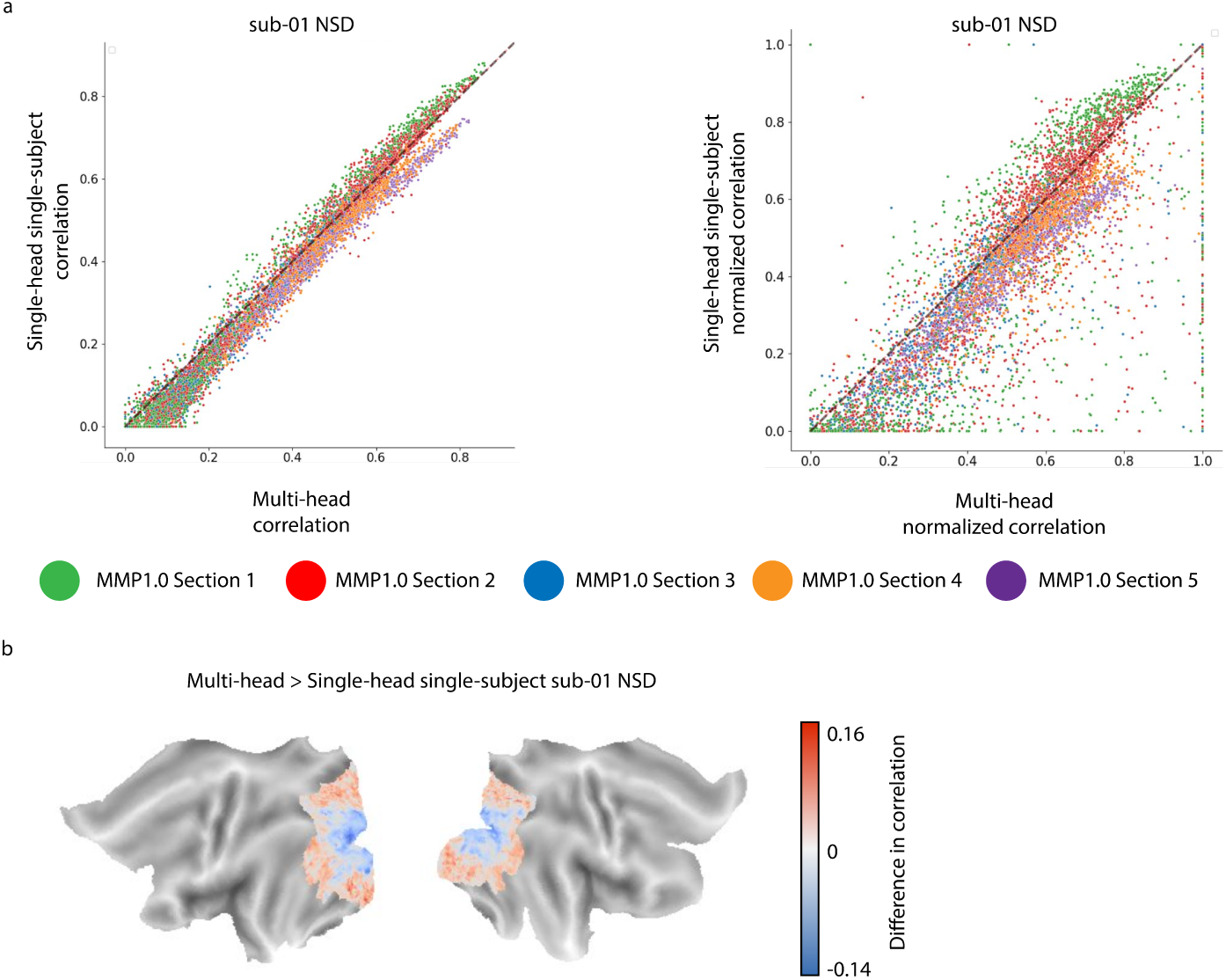
Brain-optimized model comparison scatter plot. **a** Correlation and noise normalized correlation scatter plots. Each point represents the prediction accuracy of one of the 7831 vertices from the single-head single-subject model (y-axis) trained on sub-01 NSD and the multi-head model trained on all 93 subjects (x-axis). Each model predicted sub-01 NSD’s response. Prediction accuracy is the correlation (left) or noise normalized correlation (right) between the model predictions and ground truth response. Points are color-coded according to which MMP1.0 section they belong to (MMP1.0 sections 1-5 encompasses increasingly higher-level regions of visual cortex). Points that lie above the diagonal black line mean the single-head single-subject model was the better predictor, and points that lie below the diagonal black line mean the multi-head all subject model was the better predictor. **b** Correlation difference on cortical flatmap. The flatmap shows the multi-head all subject minus the single-head single-subject (panel a, left) difference in correlation at each vertex.

**Fig. A5:**
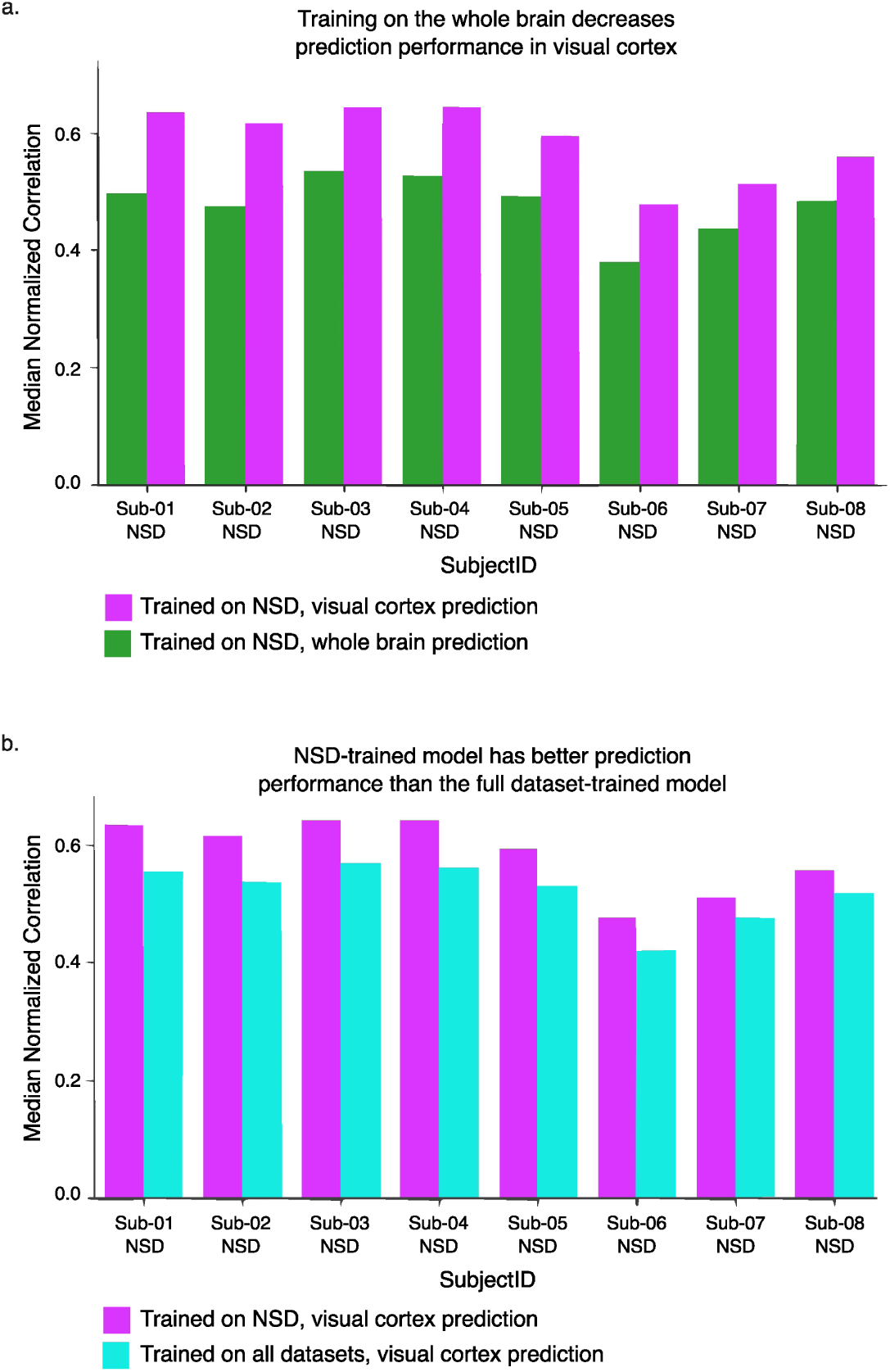
CNN8 multi-head framework training ablations. **a** Whole brain versus visual cortex training. A CNN8 multi-head model was trained on the Natural Scenes Dataset to predict the full cortex (green) and, separately, visual cortex (magenta). The prediction accuracy is the median noise normalized correlations over all of visual cortex. **b** NSD versus full dataset training. A CNN8 multi-head model was trained to predict visual cortex using brain responses from all 93 subjects in MOSAIC (cyan) and only the eight Natural Scenes Dataset subjects (magenta). The prediction accuracy is the median noise normalized correlations over all of visual cortex. The model trained on the Natural Scenes Dataset to predict visual cortex is the same in both panels (**a**) and (**b**) (magenta).

**Fig. A6:**
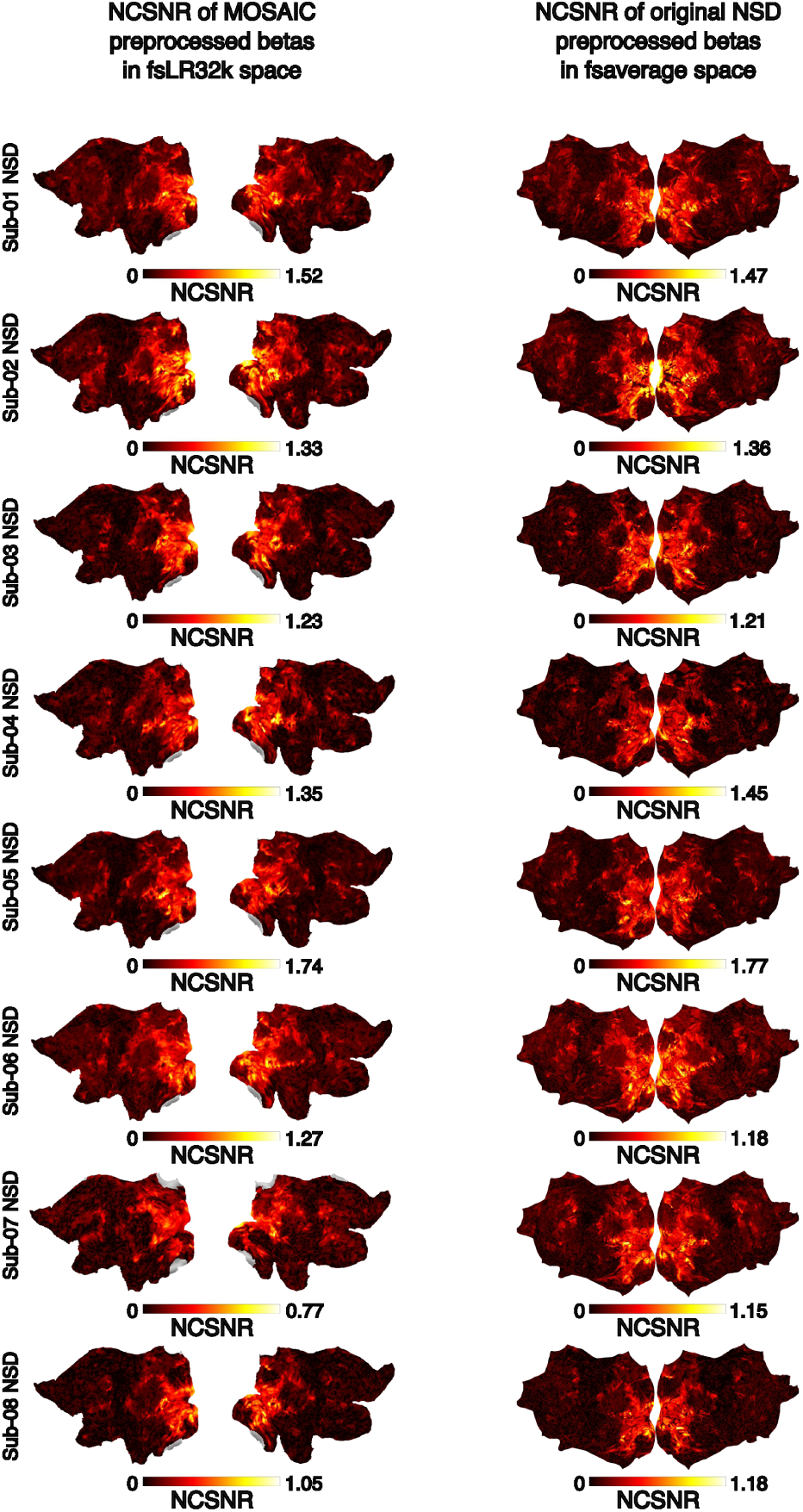
Noise ceiling signal-to-noise ratio comparison between MOSAIC and Natural Scenes Dataset (NSD) preprocessing pipelines. For each of the eight NSD subjects, the NCSNR at each vertex is plotted for the data derived from the MOSAIC preprocessing pipeline and from NSD’s original preprocessing pipeline. The NSD data used here was resampled to fsaverage space by the dataset’s original authors and most comparable to MOSAIC’s data registered to fsLR32k space. Strong spatial similarities exist between the two spaces, suggesting MOSAIC’s pipeline gives comparable results to NSD’s original pipeline.

**Fig. A7:**
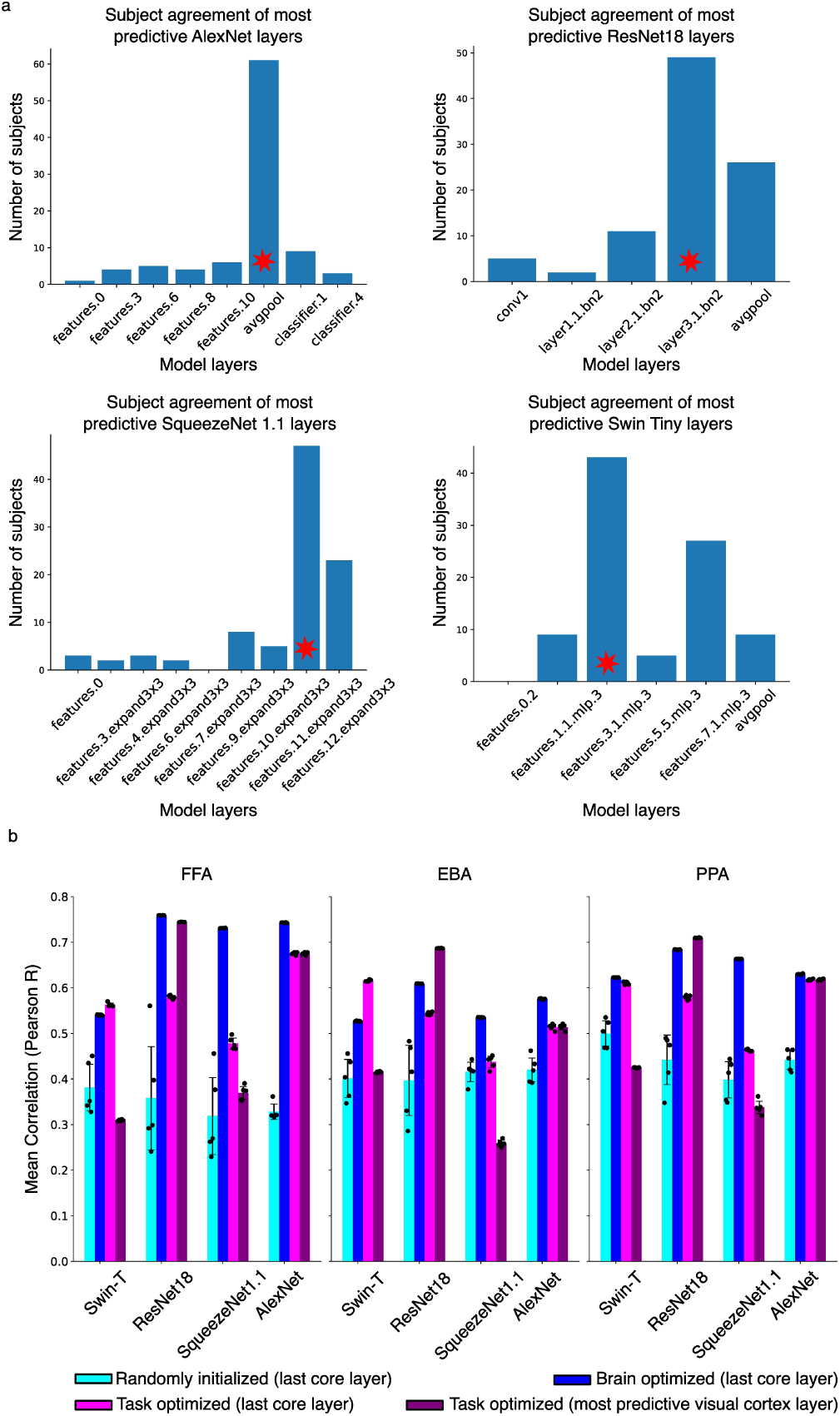
Identification of most predictive layer of task-optimized models and performance of voxelwise encoding models. **a** Model activations from the naturalistic training set images of each of MOSAIC’s 93 subjects were extracted from different layers of AlexNet, ResNet18, SqueezeNet1.1, and Swin Tiny models pre-trained on the ImageNet1k task (model architecture and weights publicly available on PyTorch). A linear model (ridge, k=5 fold cross validation, best *α* selected log-linearly between 10*^−^*^4^ and 10^8^) was trained and tested on the activations from each layer using 5-fold cross validation. The model accuracy at each layer is the median pearson correlation between the predicted and ground truth fMRI response of visual cortex (MMP1.0 sections 1-5). The histogram plots how many of the MOSAIC subjects found each layer in each model to be most predictive of visual cortex. The overall most predictive model layer of visual cortex was then selected as the layer with most subject agreement (red start). **b** Voxelwise encoding model predictions on external dataset. We extract model activations from the last core layer in models with weights optimized for brain prediction (blue), ImageNet1k task classification (magenta), and randomly initialized (cyan), and activations from the most predictive task-optimized layer (determined in **a**) (purple). A convolutional layer, followed by a linear mapping is learned between model activations and Fusiform Face Area (FFA), Parahippocampal Place Area (PPA), and Extrasriate Body Area (EBA) voxels of brain activity [105]. Error bars are standard deviations over five random seeds.

**Fig. A8:**
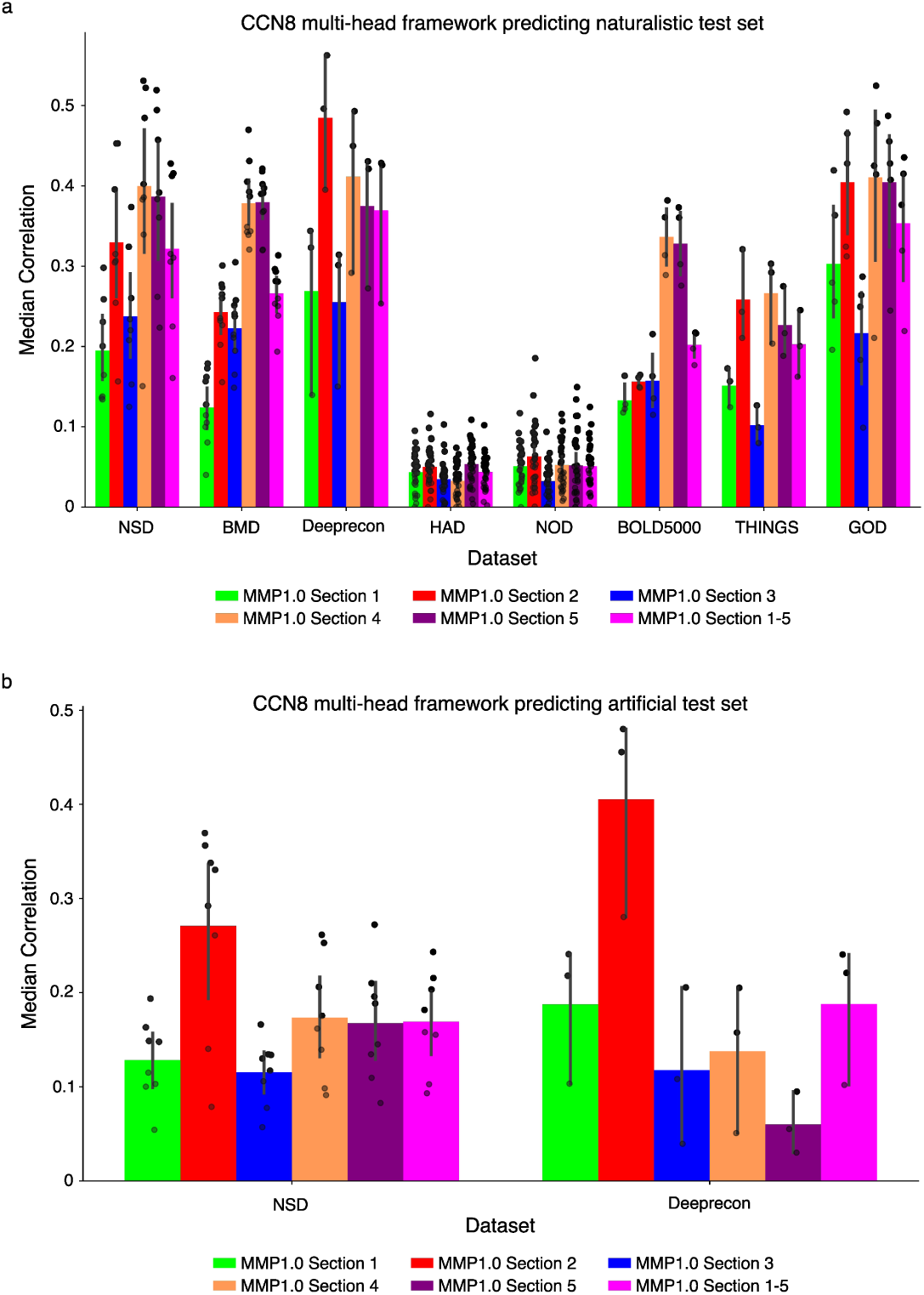
CNN8 multi-head framework prediction over all subjects. **a** Naturalistic test set prediction. The model predicts vertex-wise fMRI responses in visual cortex (MMP1.0 sections 1-5) for all 93 subjects in response to each subject’s naturalistic test set. Prediction accuracy at a given vertex is the Pearson correlation between the predicted and ground truth fMRI activity. **b** Artificial test set prediction. The model predicts vertex-wise fMRI responses in visual cortex (MMP1.0 sections 1-5) for the 8 Natural Scenes Dataset (NSD) and 3 Deeprecon subjects in response to each subject’s artificial test set. The other datasets do not have an artificial test set. Prediction accuracy at a given vertex is the Pearson correlation between the predicted and ground truth fMRI activity. In both **a** and **b** plots, each subject’s median prediction accuracy over the vertices within the individual MMP1.0 sections 1 (green), 2 (red), 3 (blue), 4 (orange), and 5 (purple) and the combined MMP1.0 sections 1-5 (pink) are plotted (black points). Error bars reflect the 95% confidence interval around the mean of the subjects within a dataset.

**Fig. A9:**
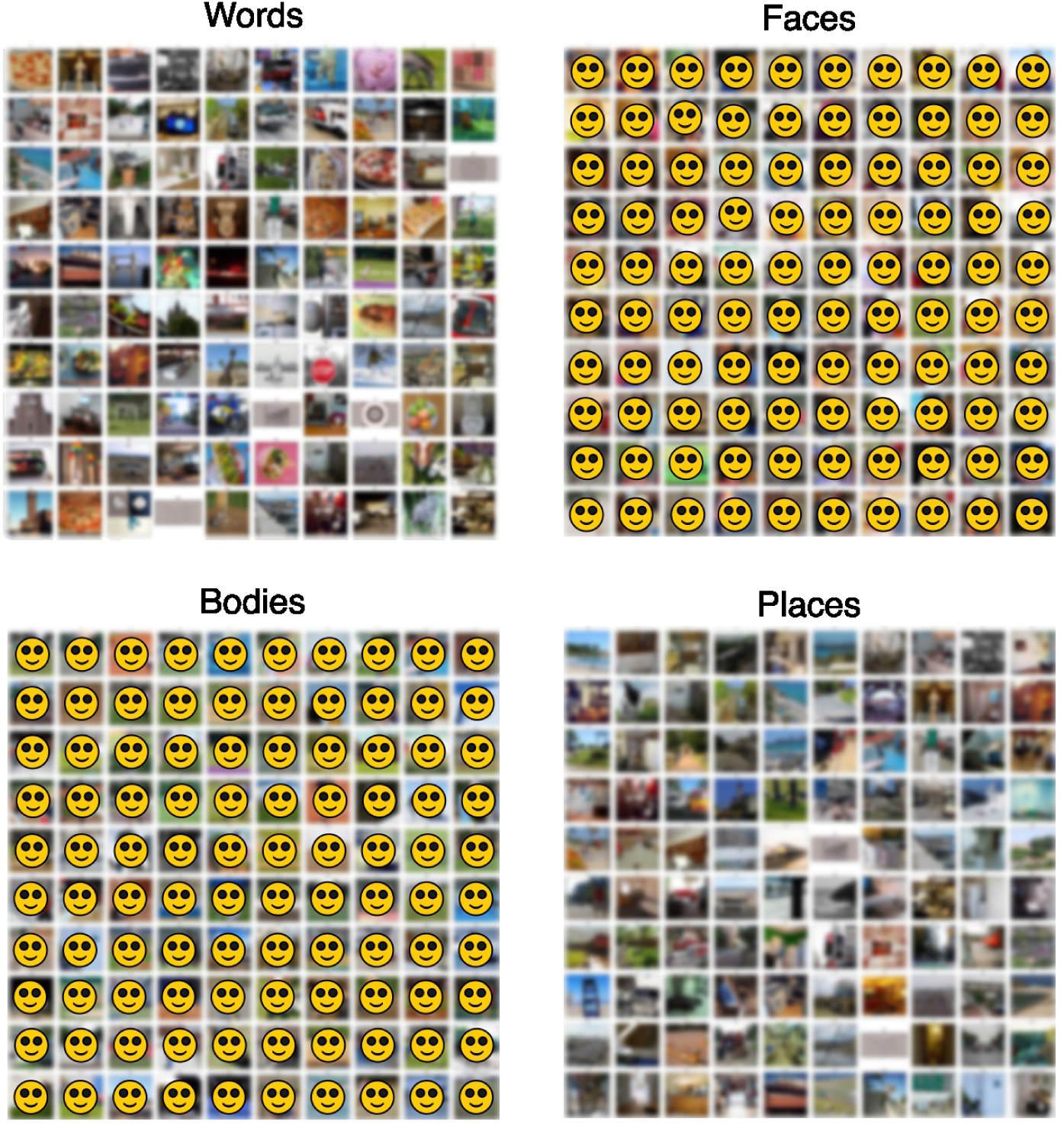
NSD sub-01 top 100 stimuli. The top 100 stimuli whose measured fMRI responses (averaged over trials) most strongly activated the vertices in the estimated contrast mask are plotted. Stimuli were ranked based on their trial-averaged fMRI response averaged across the vertices in each contrast defined by maximum F1 score. Images are blurred and identifiable faces are masked for copyright and privacy purposes. The unedited figure is available with the data release.

**Fig. A10:**
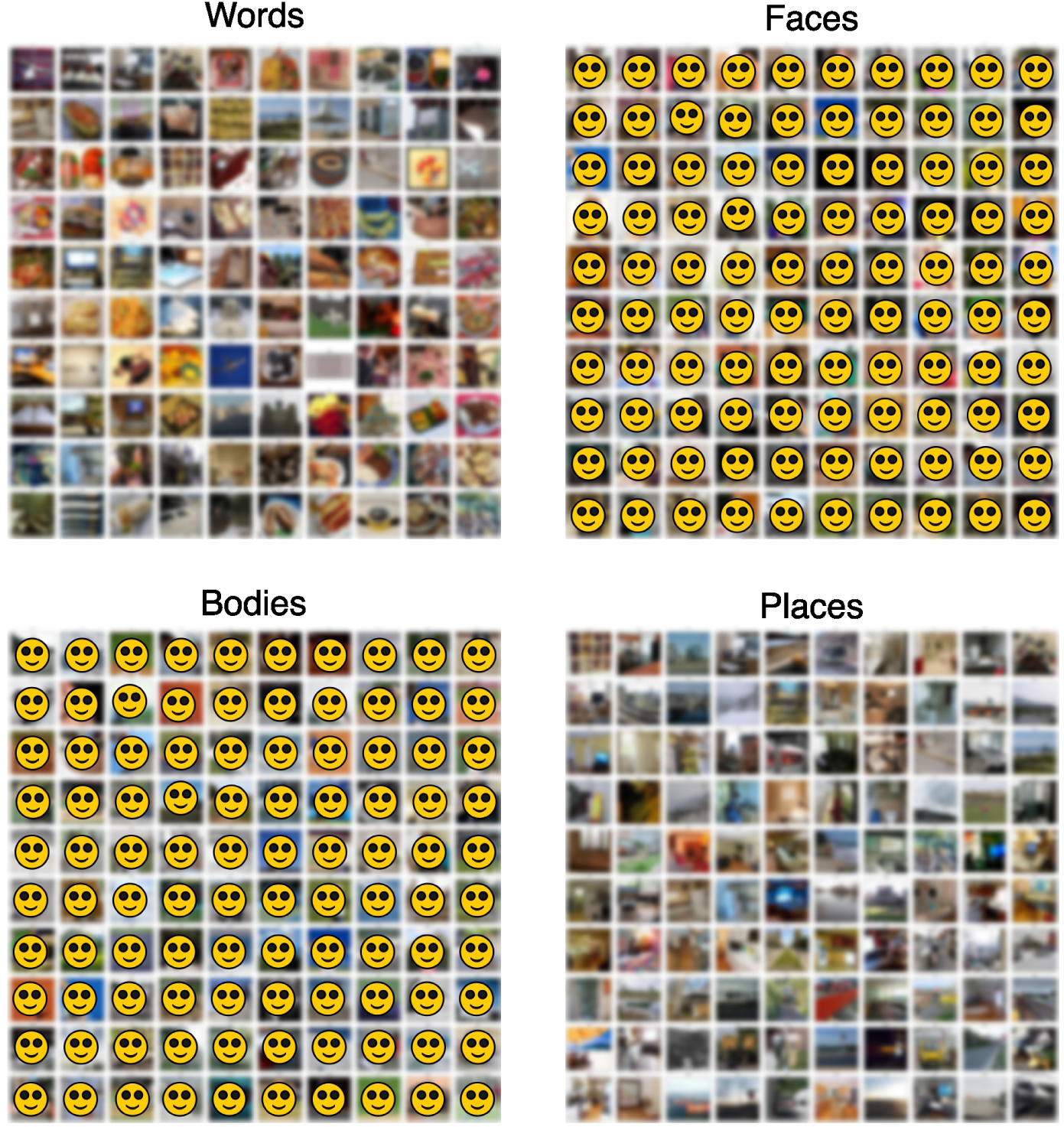
NSD sub-02 top 100 stimuli. The top 100 stimuli whose measured fMRI responses (averaged over trials) most strongly activated the vertices in the estimated contrast mask are plotted. Stimuli were ranked based on their trial-averaged fMRI response averaged across the vertices in each contrast defined by maximum F1 score. Images are blurred and identifiable faces are masked for copyright and privacy purposes. The unedited figure is available with the data release.

**Fig. A11:**
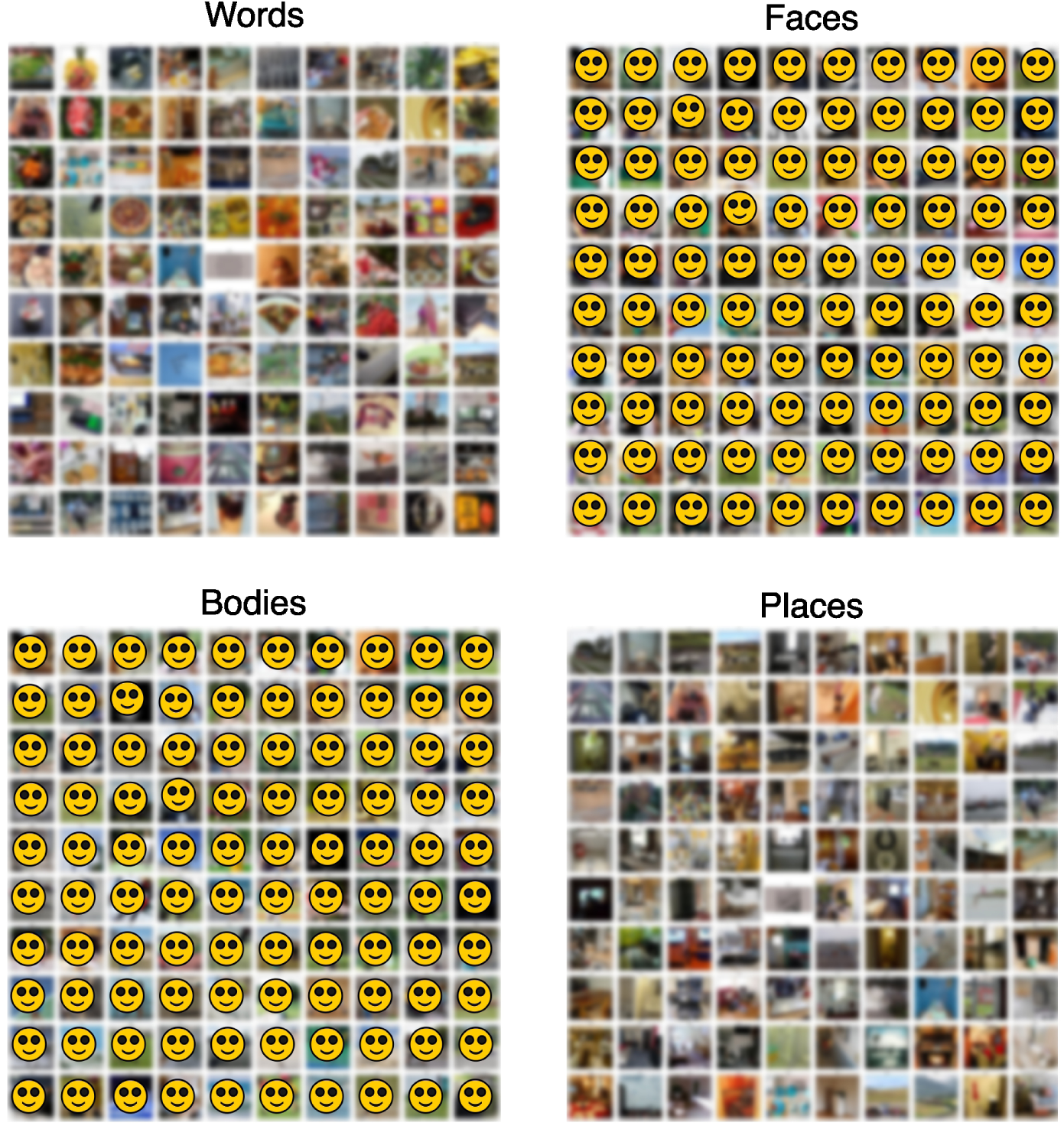
NSD sub-03 top 100 stimuli. The top 100 stimuli whose measured fMRI responses (averaged over trials) most strongly activated the vertices in the estimated contrast mask are plotted. Stimuli were ranked based on their trial-averaged fMRI response averaged across the vertices in each contrast defined by maximum F1 score. Images are blurred and identifiable faces are masked for copyright and privacy purposes. The unedited figure is available with the data release.

**Fig. A12:**
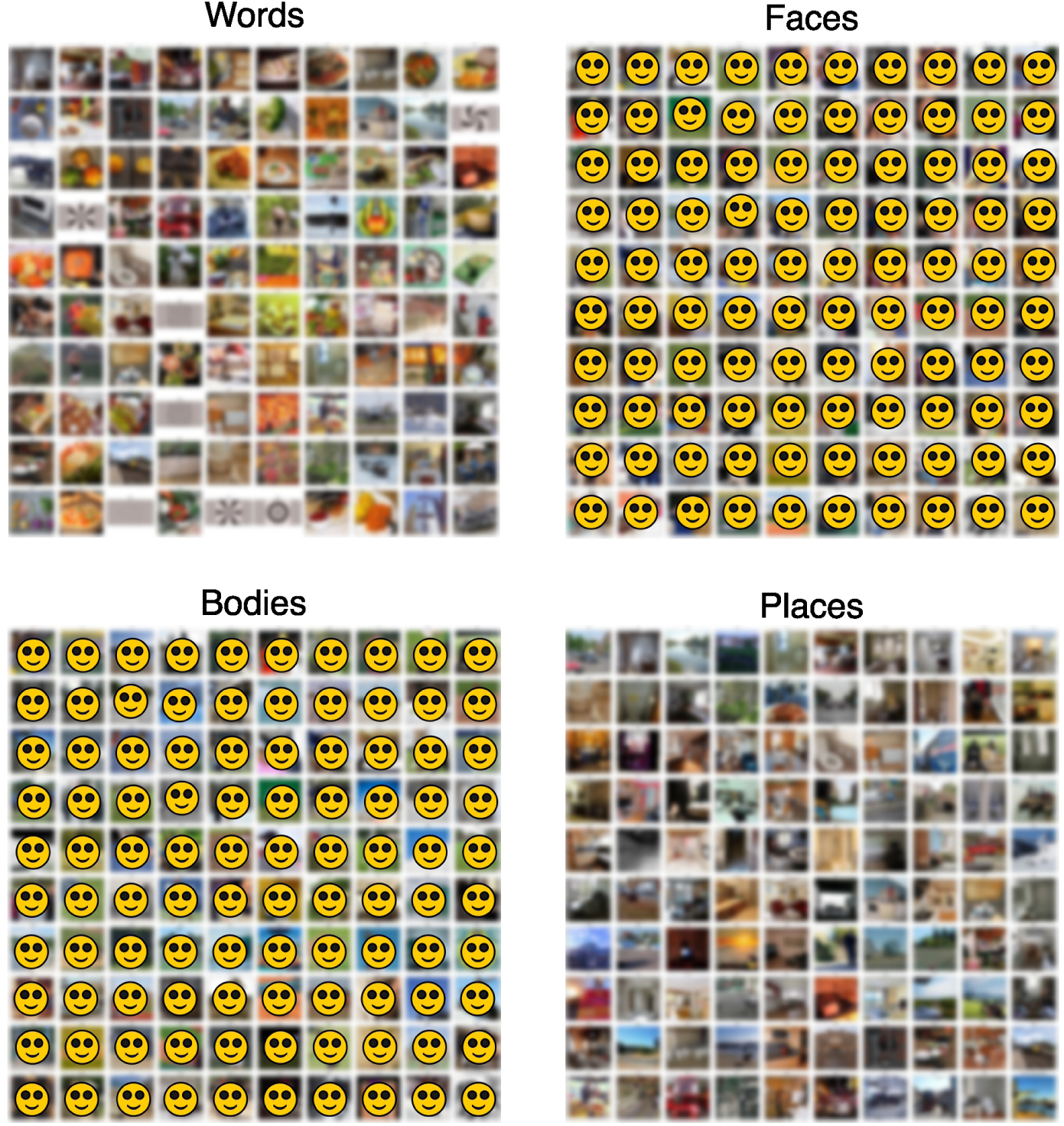
NSD sub-04 top 100 stimuli. The top 100 stimuli whose measured fMRI responses (averaged over trials) most strongly activated the vertices in the estimated contrast mask are plotted. Stimuli were ranked based on their trial-averaged fMRI response averaged across the vertices in each contrast defined by maximum F1 score. Images are blurred and identifiable faces are masked for copyright and privacy purposes. The unedited figure is available with the data release.

**Fig. A13:**
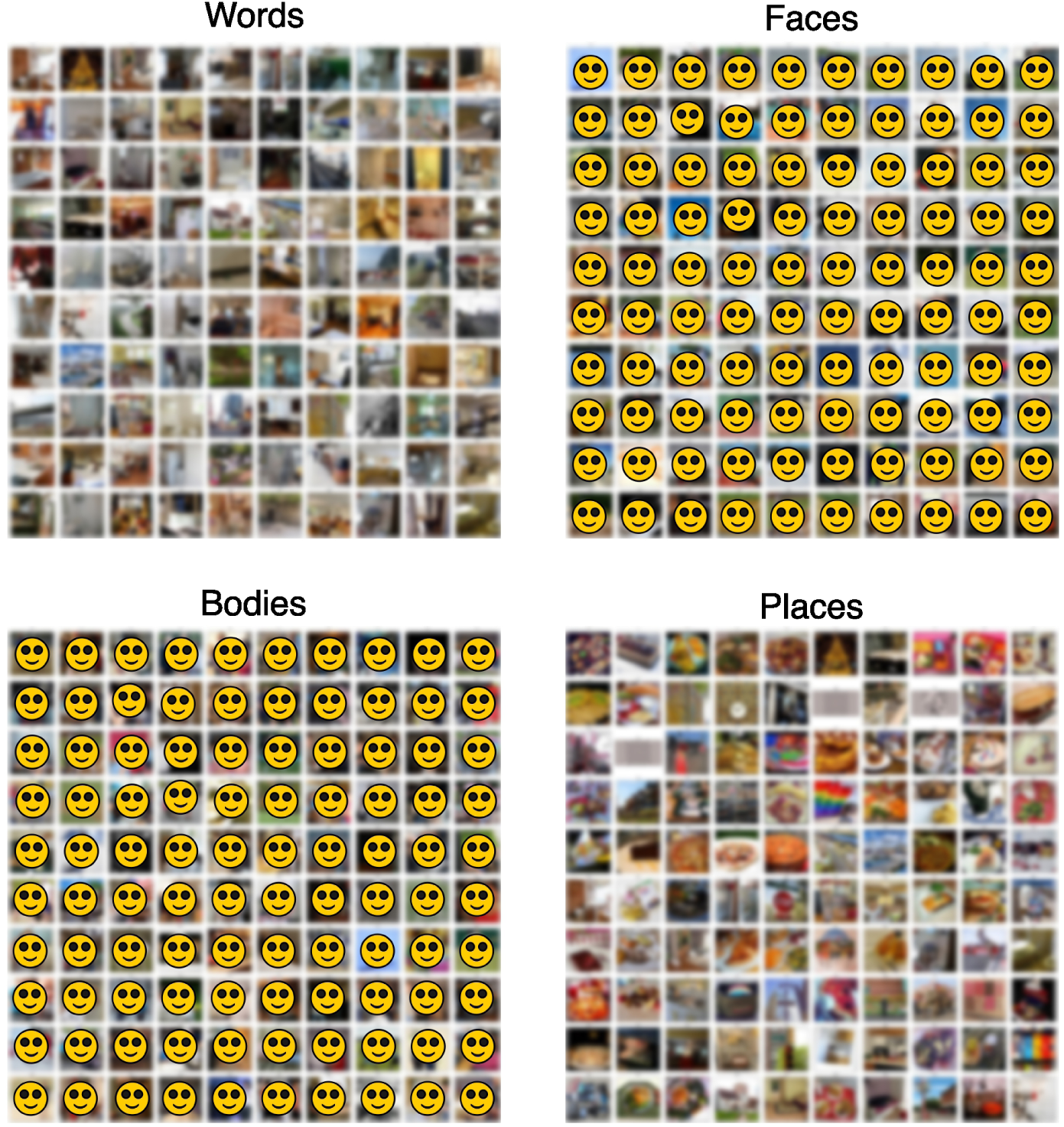
NSD sub-05 top 100 stimuli. The top 100 stimuli whose measured fMRI responses (averaged over trials) most strongly activated the vertices in the estimated contrast mask are plotted. Stimuli were ranked based on their trial-averaged fMRI response averaged across the vertices in each contrast defined by maximum F1 score. Images are blurred and identifiable faces are masked for copyright and privacy purposes. The unedited figure is available with the data release.

**Fig. A14:**
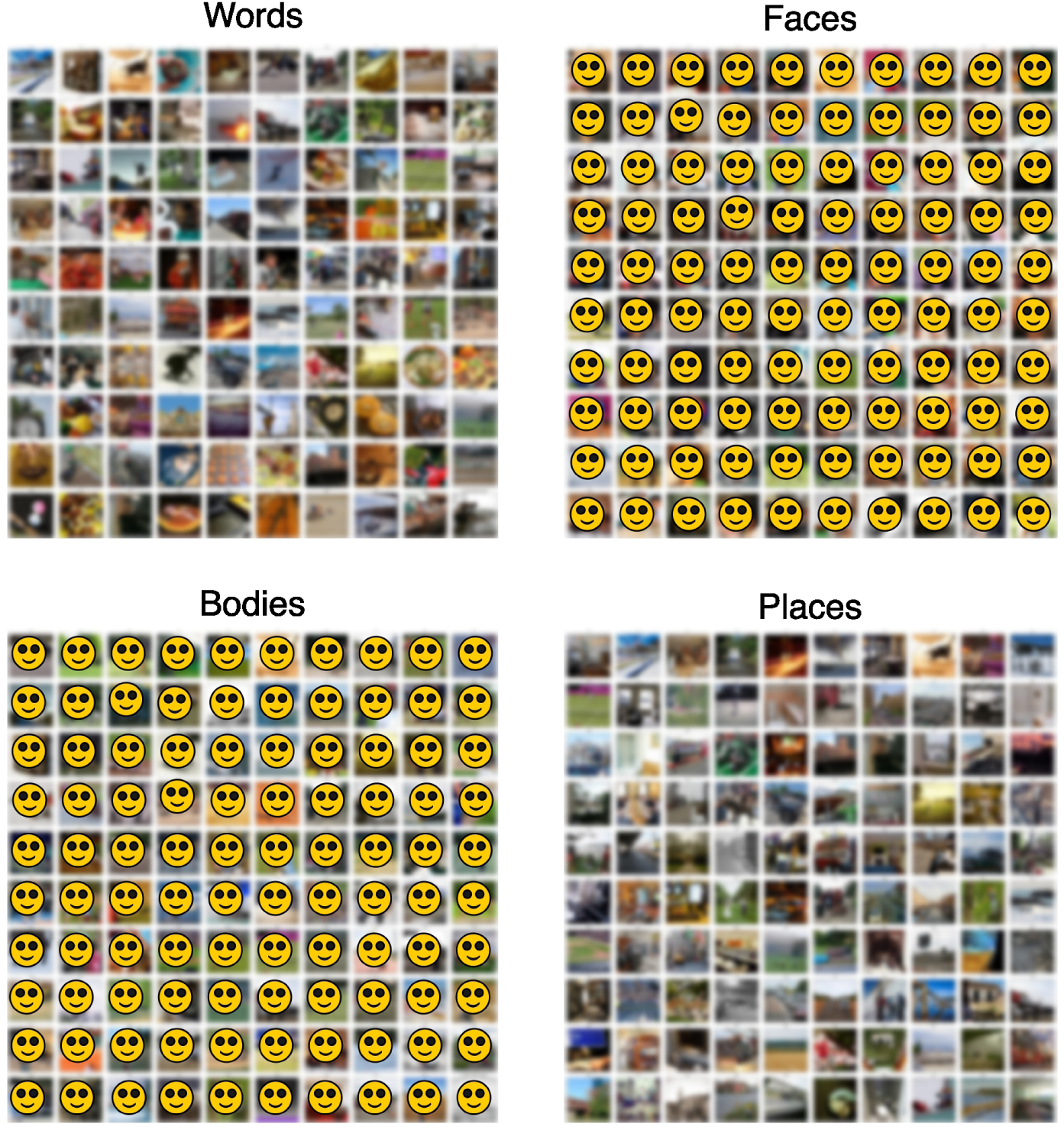
NSD sub-06 top 100 stimuli. The top 100 stimuli whose measured fMRI responses (averaged over trials) most strongly activated the vertices in the estimated contrast mask are plotted. Stimuli were ranked based on their trial-averaged fMRI response averaged across the vertices in each contrast defined by maximum F1 score. Images are blurred and identifiable faces are masked for copyright and privacy purposes. The unedited figure is available with the data release.

**Fig. A15:**
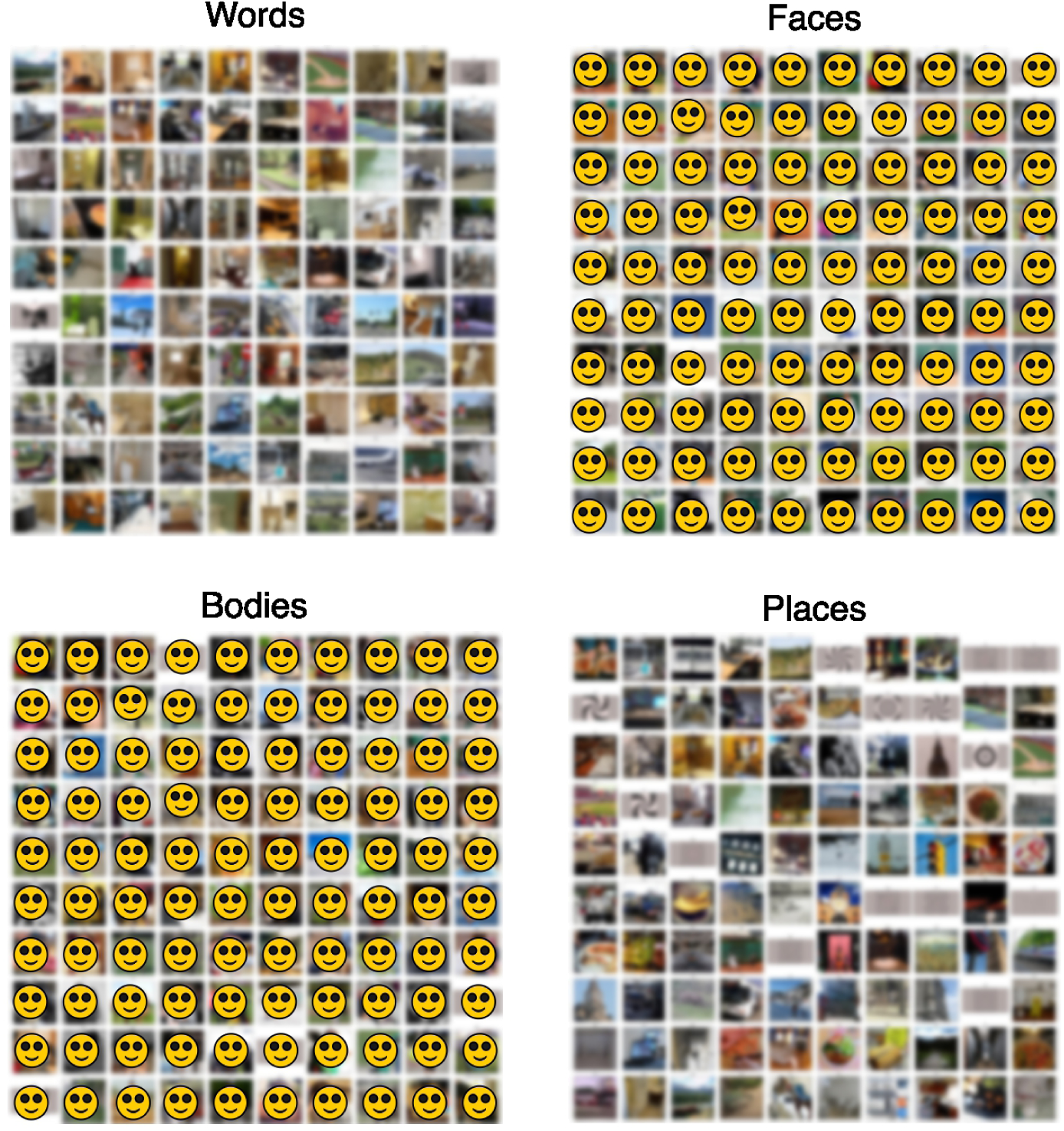
NSD sub-07 top 100 stimuli. The top 100 stimuli whose measured fMRI responses (averaged over trials) most strongly activated the vertices in the estimated contrast mask are plotted. Stimuli were ranked based on their trial-averaged fMRI response averaged across the vertices in each contrast defined by maximum F1 score. Images are blurred and identifiable faces are masked for copyright and privacy purposes. The unedited figure is available with the data release.

**Fig. A16:**
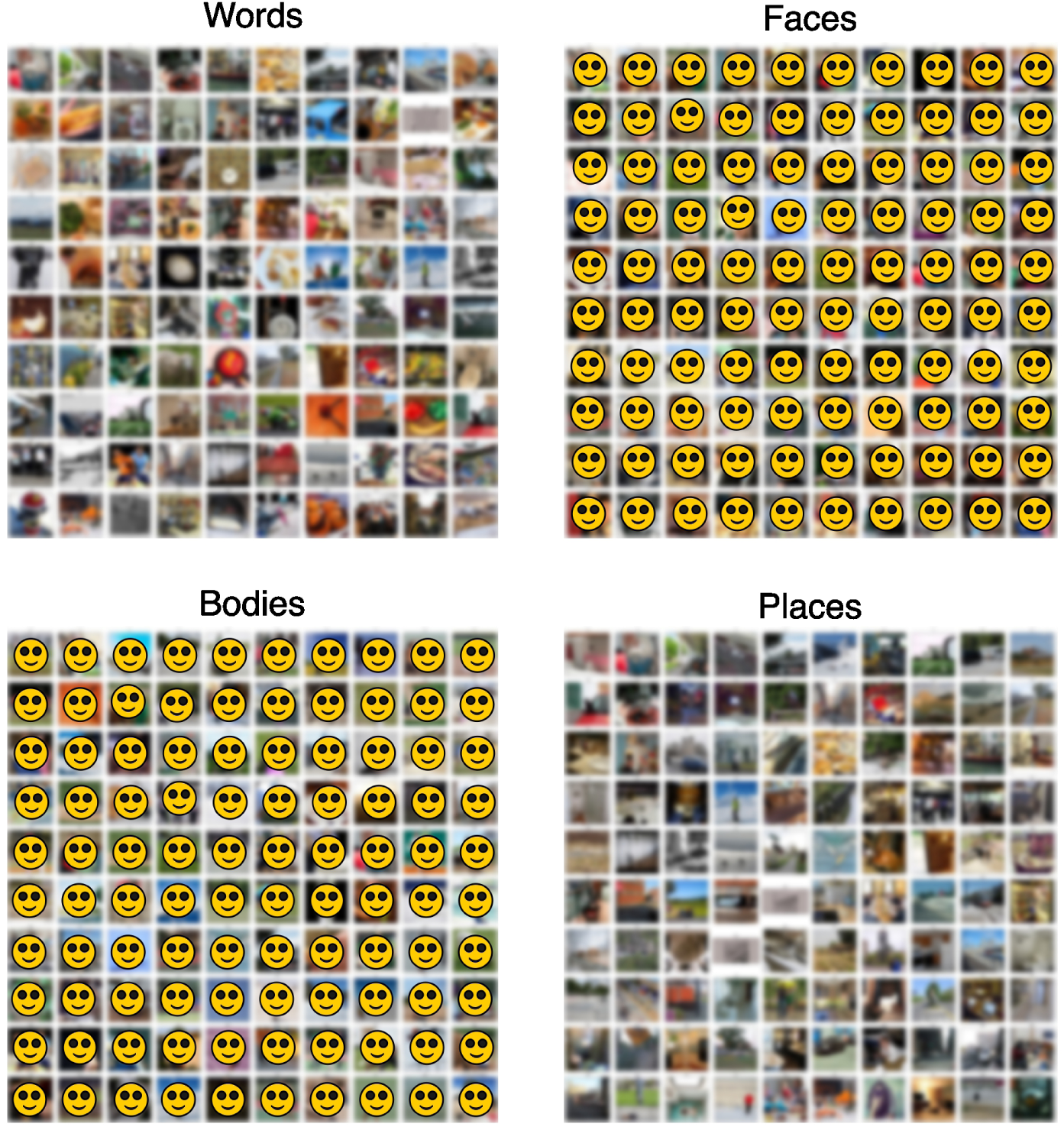
NSD sub-08 top 100 stimuli. The top 100 stimuli whose measured fMRI responses (averaged over trials) most strongly activated the vertices in the estimated contrast mask are plotted. Stimuli were ranked based on their trial-averaged fMRI response averaged across the vertices in each contrast defined by maximum F1 score. Images are blurred and identifiable faces are masked for copyright and privacy purposes. The unedited figure is available with the data release.

## Notes

https://registry.opendata.aws/mosaic/

### Competing Interest Statement

The authors have declared no competing interest.

## References

[1] Nastase, S. A., Goldstein, A. & Hasson, U. Keep it real: rethinking the primacy of experimental control in cognitive neuroscience. NeuroImage 222, 117254 (2020).

[2] Hasson, U. & Honey, C. J. Future trends in neuroimaging: Neural processes as expressed within real-life contexts. NeuroImage 62, 1272–1278 (2012).

[3] Sonkusare, S., Breakspear, M. & Guo, C. Naturalistic stimuli in neuroscience: critically acclaimed. Trends in cognitive sciences 23, 699–714 (2019).

[4] David, S. V., Vinje, W. E. & Gallant, J. L. Natural stimulus statistics alter the receptive field structure of v1 neurons. Journal of Neuroscience 24, 6991–7006 (2004).

[5] Cichy, R. M., et al. The algonauts project: A platform for communication between the sciences of biological and artificial intelligence. *arXiv preprint arXiv:1905.05675* (2019).

[6] Varoquaux, G. & Poldrack, R. A. Predictive models avoid excessive reductionism in cognitive neuroimaging. Current opinion in neurobiology 55, 1–6 (2019).

[7] Kupers, E. R., Knapen, T., Merriam, E. P. & Kay, K. N. Principles of intensive human neuroimaging. Trends in Neurosciences (2024).

[8] Yarkoni, T. & Westfall, J. Choosing prediction over explanation in psychology: Lessons from machine learning. Perspectives on Psychological Science 12, 1100– 1122 (2017).

[9] Van Essen, D. C. et al. The wu-minn human connectome project: an overview. Neuroimage 80, 62–79 (2013).

[10] Conwell, C., Prince, J. S., Kay, K. N., Alvarez, G. A. & Konkle, T. A large-scale examination of inductive biases shaping high-level visual representation in brains and machines. Nature communications 15, 9383 (2024).

[11] Allen, E. J. et al. A massive 7t fmri dataset to bridge cognitive neuroscience and artificial intelligence. Nature neuroscience 25, 116–126 (2022).

[12] Khosla, M., Jamison, K., Kuceyeski, A. & Sabuncu, M. Characterizing the ventral visual stream with response-optimized neural encoding models. Advances in Neural Information Processing Systems 35, 9389–9402 (2022).

[13] Banville, H., Benchetrit, Y., d’Ascoli, S., Rapin, J. & King, J.-R. Scaling laws for decoding images from brain activity. *arXiv preprint arXiv:2501.15322* (2025).

[14] Naselaris, T., Allen, E. & Kay, K. Extensive sampling for complete models of individual brains. Current Opinion in Behavioral Sciences 40, 45–51 (2021).

[15] Lahner, B. et al. Modeling short visual events through the bold moments video fmri dataset and metadata. Nature communications 15, 6241 (2024).

[16] Lingnau, A. & Downing, P. E. The lateral occipitotemporal cortex in action. Trends in cognitive sciences 19, 268–277 (2015).

[17] Yildirim, I., Wu, J., Kanwisher, N. & Tenenbaum, J. An integrative computational architecture for object-driven cortex. Current opinion in neurobiology 55, 73–81 (2019).

[18] Hasson, U. et al. Neurocinematics: The neuroscience of film. Projections 2, 1–26 (2008).

[19] Hasson, U., Furman, O., Clark, D., Dudai, Y. & Davachi, L. Enhanced inter-subject correlations during movie viewing correlate with successful episodic encoding. Neuron 57, 452–462 (2008).

[20] Boyle, J. A., et al. The courtois project on neuronal modelling-first data release (2020).

[21] Hanke, M. et al. A studyforrest extension, simultaneous fmri and eye gaze recordings during prolonged natural stimulation. Scientific data 3, 1–15 (2016).

[22] Eickhoff, S., Nichols, T. E., Van Horn, J. D. & Turner, J. A. Sharing the wealth: Neuroimaging data repositories. Neuroimage 124, 1065 (2016).

[23] Gorgolewski, K. J. et al. The brain imaging data structure, a format for organizing and describing outputs of neuroimaging experiments. Scientific data 3, 1–9 (2016).

[24] Markiewicz, C. J. et al. The openneuro resource for sharing of neuroscience data. Elife 10, e71774 (2021).

[25] Dubois, J. & Adolphs, R. Building a science of individual differences from fmri. Trends in cognitive sciences 20, 425–443 (2016).

[26] Bennett, C. M. & Miller, M. B. fmri reliability: Influences of task and experimental design. Cognitive, Affective, & Behavioral Neuroscience 13, 690–702 (2013).

[27] Matthews, P. M., Honey, G. D. & Bullmore, E. T. Applications of fmri in translational medicine and clinical practice. Nature Reviews Neuroscience 7, 732–744 (2006).

[28] D’Esposito, M., Deouell, L. Y. & Gazzaley, A. Alterations in the bold fmri signal with ageing and disease: a challenge for neuroimaging. Nature Reviews Neuroscience 4, 863–872 (2003).

[29] Friedman, L., Glover, G. H., Krenz, D., Magnotta, V. & Birn, T. F. Reducing inter-scanner variability of activation in a multicenter fmri study: role of smoothness equalization. Neuroimage 32, 1656–1668 (2006).

[30] Thirion, B. et al. Analysis of a large fmri cohort: Statistical and methodological issues for group analyses. Neuroimage 35, 105–120 (2007).

[31] Sun, J. et al. Contrast, attend and diffuse to decode high-resolution images from brain activities. Advances in Neural Information Processing Systems 36, 12332–12348 (2023).

[32] Caro, J. O. et al. Brainlm: A foundation model for brain activity recordings. bioRxiv 2023–09 (2023).

[33] Dong, Z. et al. Brain-jepa: Brain dynamics foundation model with gradient positioning and spatiotemporal masking. Advances in Neural Information Processing Systems 37, 86048–86073 (2025).

[34] Mensch, A., Mairal, J., Thirion, B. & Varoquaux, G. Extracting representations of cognition across neuroimaging studies improves brain decoding. PLoS computational biology 17, e1008795 (2021).

[35] Thomas, A., Ŕe, C. & Poldrack, R. Self-supervised learning of brain dynamics from broad neuroimaging data. Advances in neural information processing systems 35, 21255–21269 (2022).

[36] Fosco, C., et al. Brain netflix: Scaling data to reconstruct videos from brain signals, 457–474 (Springer, 2025).

[37] Turner, B. O., Paul, E. J., Miller, M. B. & Barbey, A. K. Small sample sizes reduce the replicability of task-based fmri studies. Communications biology 1, 62 (2018).

[38] Cremers, H. R., Wager, T. D. & Yarkoni, T. The relation between statistical power and inference in fmri. PloS one 12, e0184923 (2017).

[39] Murphy, K. & Garavan, H. An empirical investigation into the number of subjects required for an event-related fmri study. Neuroimage 22, 879–885 (2004).

[40] Esteban, O. et al. fmriprep: a robust preprocessing pipeline for functional mri. Nature methods 16, 111–116 (2019).

[41] Prince, J. S. et al. Improving the accuracy of single-trial fmri response estimates using glmsingle. Elife 11, e77599 (2022).

[42] Fu, S. et al. Dreamsim: Learning new dimensions of human visual similarity using synthetic data. Advances in Neural Information Processing Systems 36 (2024).

[43] Glasser, M. F. et al. A multi-modal parcellation of human cerebral cortex. Nature 536, 171–178 (2016).

[44] Aurich, N. K., Alves Filho, J. O., Marques da Silva, A. M. & Franco, A. R. Evaluating the reliability of different preprocessing steps to estimate graph theoretical measures in resting state fmri data. Frontiers in neuroscience 9, 48 (2015).

[45] Kriegeskorte, N. & Douglas, P. K. Interpreting encoding and decoding models. Current opinion in neurobiology 55, 167–179 (2019).

[46] Horikawa, T. & Kamitani, Y. Generic decoding of seen and imagined objects using hierarchical visual features. Nature communications 8, 15037 (2017).

[47] Shen, G., Horikawa, T., Majima, K. & Kamitani, Y. Deep image reconstruction from human brain activity. PLoS computational biology 15, e1006633 (2019).

[48] Hebart, M. N. et al. Things-data, a multimodal collection of large-scale datasets for investigating object representations in human brain and behavior. Elife 12, e82580 (2023).

[49] Kriegeskorte, N., Mur, M. & Bandettini, P. A. Representational similarity analysis-connecting the branches of systems neuroscience. Frontiers in systems neuroscience 2, 249 (2008).

[50] Haxby, J. V., Guntupalli, J. S., Nastase, S. A. & Feilong, M. Hyperalignment: Modeling shared information encoded in idiosyncratic cortical topographies. elife 9, e56601 (2020).

[51] Mazziotta, J. C. et al. A probabilistic atlas of the human brain: theory and rationale for its development. Neuroimage 2, 89–101 (1995).

[52] Glasser, M. F. et al. The minimal preprocessing pipelines for the human connectome project. Neuroimage 80, 105–124 (2013).

[53] Robinson, E. C. et al. Msm: a new flexible framework for multimodal surface matching. Neuroimage 100, 414–426 (2014).

[54] Robinson, E. C. et al. Multimodal surface matching with higher-order smoothness constraints. Neuroimage 167, 453–465 (2018).

[55] Van Essen, D. C., Glasser, M. F., Dierker, D. L., Harwell, J. & Coalson, T. Parcellations and hemispheric asymmetries of human cerebral cortex analyzed on surface-based atlases. Cerebral cortex 22, 2241–2262 (2012).

[56] Coalson, T. S., Van Essen, D. C. & Glasser, M. F. The impact of traditional neuroimaging methods on the spatial localization of cortical areas. Proceedings of the National Academy of Sciences 115, E6356–E6365 (2018).

[57] Brodoehl, S., Gaser, C., Dahnke, R., Witte, O. W. & Klingner, C. M. Surface-based analysis increases the specificity of cortical activation patterns and connectivity results. Scientific reports 10, 5737 (2020).

[58] Jeganathan, J. et al. Spurious correlations in surface-based functional brain imaging. Imaging Neuroscience (2025).

[59] Smith, S. M. et al. Resting-state fmri in the human connectome project. Neuroimage 80, 144–168 (2013).

[60] Satterthwaite, T. D. et al. An improved framework for confound regression and filtering for control of motion artifact in the preprocessing of resting-state functional connectivity data. Neuroimage 64, 240–256 (2013).

[61] Chen, Z., Qing, J., Xiang, T., Yue, W. L. & Zhou, J. H. Seeing beyond the brain: Conditional diffusion model with sparse masked modeling for vision decoding, 22710–22720 (2023).

[62] Chen, Z., Qing, J. & Zhou, J. H. Cinematic mindscapes: High-quality video reconstruction from brain activity. Advances in Neural Information Processing Systems 36 (2024).

[63] Fortin, J.-P. et al. Harmonization of multi-site diffusion tensor imaging data. Neuroimage 161, 149–170 (2017).

[64] Costafreda, S. G. et al. Multisite fmri reproducibility of a motor task using identical mr systems. Journal of Magnetic Resonance Imaging: An Official Journal of the International Society for Magnetic Resonance in Medicine 26, 1122–1126 (2007).

[65] Suckling, J. et al. Components of variance in a multicentre functional mri study and implications for calculation of statistical power. Human Brain Mapping 29, 1111–1122 (2008).

[66] Zou, K. H. et al. Reproducibility of functional mr imaging: preliminary results of prospective multi-institutional study performed by biomedical informatics research network. Radiology 237, 781–789 (2005).

[67] Brown, G. G. et al. Multisite reliability of cognitive bold data. Neuroimage 54, 2163–2175 (2011).

[68] Saygin, Z. M. et al. Connectivity precedes function in the development of the visual word form area. Nature neuroscience 19, 1250–1255 (2016).

[69] Willems, R. M., Peelen, M. V. & Hagoort, P. Cerebral lateralization of face-selective and body-selective visual areas depends on handedness. Cerebral cortex 20, 1719–1725 (2010).

[70] Van Horn, J. D., Grafton, S. T. & Miller, M. B. Individual variability in brain activity: a nuisance or an opportunity? Brain imaging and behavior 2, 327–334 (2008).

[71] Julian, J. B., Fedorenko, E., Webster, J. & Kanwisher, N. An algorithmic method for functionally defining regions of interest in the ventral visual pathway. Neuroimage 60, 2357–2364 (2012).

[72] Nieto-Castanon, A., Ghosh, S. S., Tourville, J. A. & Guenther, F. H. Region of interest based analysis of functional imaging data. Neuroimage 19, 1303–1316 (2003).

[73] Naselaris, T., Kay, K. N., Nishimoto, S. & Gallant, J. L. Encoding and decoding in fmri. Neuroimage 56, 400–410 (2011).

[74] Tang, J., LeBel, A., Jain, S. & Huth, A. G. Semantic reconstruction of continuous language from non-invasive brain recordings. Nature Neuroscience 26, 858–866 (2023).

[75] Kneeland, R., et al. Nsd-imagery: A benchmark dataset for extending fmri vision decoding methods to mental imagery, 28852–28862 (2025).

[76] Diedrichsen, J., Wiestler, T. & Krakauer, J. W. Two distinct ipsilateral cortical representations for individuated finger movements. Cerebral Cortex 23, 1362– 1377 (2013).

[77] Roth, J. & Hebart, M. N. How to sample the world for understanding the visual system (2025).

[78] Shirakawa, K. et al. Spurious reconstruction from brain activity: The thin line between reconstruction, classification, and hallucination. Journal of Vision 24, 321–321 (2024).

[79] Wang, A. Y., Kay, K., Naselaris, T., Tarr, M. J. & Wehbe, L. Better models of human high-level visual cortex emerge from natural language supervision with a large and diverse dataset. Nature Machine Intelligence 5, 1415–1426 (2023).

[80] Miller, K. L. et al. Multimodal population brain imaging in the uk biobank prospective epidemiological study. Nature neuroscience 19, 1523–1536 (2016).

[81] Taylor, J. R. et al. The cambridge centre for ageing and neuroscience (cam-can) data repository: Structural and functional mri, meg, and cognitive data from a cross-sectional adult lifespan sample. neuroimage 144, 262–269 (2017).

[82] Casey, B. J. et al. The adolescent brain cognitive development (abcd) study: imaging acquisition across 21 sites. Developmental cognitive neuroscience 32, 43–54 (2018).

[83] Zhang, Y. et al. Video instruction tuning with synthetic data. arXiv preprint arXiv:2410.02713 (2024).

[84] Bolya, D. et al. Perception encoder: The best visual embeddings are not at the output of the network. arXiv preprint arXiv:2504.13181 (2025).

[85] Schuhmann, C. et al. Laion-5b: An open large-scale dataset for training next generation image-text models. Advances in neural information processing systems 35, 25278–25294 (2022).

[86] Yamins, D. L., Hong, H., Cadieu, C. & DiCarlo, J. J. Hierarchical modular optimization of convolutional networks achieves representations similar to macaque it and human ventral stream. Advances in neural information processing systems 26 (2013).

[87] Schrimpf, M. et al. Integrative benchmarking to advance neurally mechanistic models of human intelligence. Neuron 108, 413–423 (2020).

[88] Cichy, R. M., Khosla, A., Pantazis, D., Torralba, A. & Oliva, A. Comparison of deep neural networks to spatio-temporal cortical dynamics of human visual object recognition reveals hierarchical correspondence. Scientific reports 6, 27755 (2016).

[89] Gallant, J. L., Nishimoto, S., Naselaris, T. & Wu, M. System identification, encoding models, and decoding models: a powerful new approach to fmri research. Visual population codes: Toward a common multivariate framework for cell recording and functional imaging 163–188 (2012).

[90] Wu, M. C.-K., David, S. V. & Gallant, J. L. Complete functional characterization of sensory neurons by system identification. Annu. Rev. Neurosci. 29, 477–505 (2006).

[91] Seeliger, K. et al. End-to-end neural system identification with neural information flow. PLOS Computational Biology 17, e1008558 (2021).

[92] St-Yves, G., Allen, E. J., Wu, Y., Kay, K. & Naselaris, T. Brain-optimized deep neural network models of human visual areas learn non-hierarchical representations. Nature communications 14, 3329 (2023).

[93] Olivetti, E., Greiner, S. & Avesani, P. Adhd diagnosis from multiple data sources with batch effects. Frontiers in systems neuroscience 6, 70 (2012).

[94] Yu, M. et al. Statistical harmonization corrects site effects in functional connectivity measurements from multi-site fmri data. Human brain mapping 39, 4213–4227 (2018).

[95] Klindt, D., Ecker, A. S., Euler, T. & Bethge, M. Neural system identification for large populations separating “what” and “where”. Advances in neural information processing systems 30 (2017).

[96] Beliy, R., Wasserman, N., Zalcher, A. & Irani, M. The wisdom of a crowd of brains: A universal brain encoder. *arXiv preprint arXiv:2406.12179* (2024).

[97] Krizhevsky, A., Sutskever, I. & Hinton, G. E. Imagenet classification with deep convolutional neural networks. Advances in neural information processing systems 25 (2012).

[98] Iandola, F. N., et al. Squeezenet: Alexnet-level accuracy with 50x fewer parameters and¡ 0.5 mb model size. *arXiv preprint arXiv:1602.07360* (2016).

[99] He, K., Zhang, X., Ren, S. & Sun, J. Deep residual learning for image recognition, 770–778 (2016).

[100] Liu, Z., et al. Swin transformer: Hierarchical vision transformer using shifted windows, 10012–10022 (2021).

[101] Kietzmann, T. C. et al. Recurrence is required to capture the representational dynamics of the human visual system. Proceedings of the National Academy of Sciences 116, 21854–21863 (2019).

[102] Spoerer, C. J., McClure, P. & Kriegeskorte, N. Recurrent convolutional neural networks: a better model of biological object recognition. Frontiers in psychology 8, 1551 (2017).

[103] Kubilius, J. et al. Brain-like object recognition with high-performing shallow recurrent anns. Advances in neural information processing systems 32 (2019).

[104] Koivisto, M., Railo, H., Revonsuo, A., Vanni, S. & Salminen-Vaparanta, N. Recurrent processing in v1/v2 contributes to categorization of natural scenes. Journal of Neuroscience 31, 2488–2492 (2011).

[105] Ratan Murty, N. A., Bashivan, P., Abate, A., DiCarlo, J. J. & Kanwisher, N. Computational models of category-selective brain regions enable high-throughput tests of selectivity. Nature communications 12, 5540 (2021).

[106] Stigliani, A., Weiner, K. S. & Grill-Spector, K. Temporal processing capacity in high-level visual cortex is domain specific. Journal of Neuroscience 35, 12412– 12424 (2015).

[107] [107] Torralba, A. & Efros, A. A. Unbiased look at dataset bias, 1521–1528 (IEEE, 2011).

[108] Zeng, B., Yin, Y. & Liu, Z. Understanding bias in large-scale visual datasets. *arXiv preprint arXiv:2412.01876* (2024).

[109] Liu, Z. & He, K. A decade’s battle on dataset bias: Are we there yet? *arXiv preprint arXiv:2403*.*08632* (2024).

[110] Oliva, A. & Torralba, A. The role of context in object recognition. Trends in cognitive sciences 11, 520–527 (2007).

[111] Hebart, M. N. et al. Things: A database of 1,854 object concepts and more than 26,000 naturalistic object images. PloS one 14, e0223792 (2019).

[112] Arcaro, M. J., Ponce, C. & Livingstone, M. The neurons that mistook a hat for a face. Elife 9, e53798 (2020).

[113] Caron, M., et al. Emerging properties in self-supervised vision transformers, 9650–9660 (2021).

[114] Zhai, X., Kolesnikov, A., Houlsby, N. & Beyer, L. Scaling vision transformers, 12104–12113 (2022).

[115] Radford, A. et al. Learning transferable visual models from natural language supervision, 8748–8763 (PmLR, 2021).

[116] Sun, C., Shrivastava, A., Singh, S. & Gupta, A. Revisiting unreasonable effectiveness of data in deep learning era, 843–852 (2017).

[117] Kriegeskorte, N. Deep neural networks: a new framework for modeling biological vision and brain information processing. Annual review of vision science 1, 417–446 (2015).

[118] Muttenthaler, L., Dippel, J., Linhardt, L., Vandermeulen, R. A. & Kornblith, S. Human alignment of neural network representations. *arXiv preprint arXiv:2211.01201* (2022).

[119] Fel, T., Rodriguez Rodriguez, I. F., Linsley, D. & Serre, T. Harmonizing the object recognition strategies of deep neural networks with humans. Advances in neural information processing systems 35, 9432–9446 (2022).

[120] Pant, N., Rodriguez, I. F., Beniwal, A., Warren, S. & Serre, T. Hmax strikes back: Self-supervised learning of human-like scale invariant representations (2024).

[121] Konkle, T. & Alvarez, G. A. A self-supervised domain-general learning framework for human ventral stream representation. Nature communications 13, 491 (2022).

[122] Huh, M., Cheung, B., Wang, T. & Isola, P. The platonic representation hypothesis. *arXiv preprint arXiv:2405.07987* (2024).

[123] Gifford, A. T., Jastrzebowska, M. A., Singer, J. J. & Cichy, R. M. In silico discovery of representational relationships across visual cortex. Nature Human Behaviour 1–20 (2025).

[124] Chang, N. et al. Bold5000, a public fmri dataset while viewing 5000 visual images. Scientific data 6, 49 (2019).

[125] Lin, T.-Y., et al. Microsoft coco: Common objects in context, 740–755 (Springer, 2014).

[126] Xiao, J., Hays, J., Ehinger, K. A., Oliva, A. & Torralba, A. Sun database: Large-scale scene recognition from abbey to zoo, 3485–3492 (IEEE, 2010).

[127] Deng, J., et al. Imagenet: A large-scale hierarchical image database, 248–255 (Ieee, 2009).

[128] Monfort, M. et al. Multi-moments in time: Learning and interpreting models for multi-action video understanding. IEEE Transactions on Pattern Analysis and Machine Intelligence 44, 9434–9445 (2021).

[129] Monfort, M. et al. Moments in time dataset: one million videos for event understanding. IEEE transactions on pattern analysis and machine intelligence 42, 502–508 (2019).

[130] Zhou, M. et al. A large-scale fmri dataset for human action recognition. Scientific Data 10, 415 (2023).

[131] Zhao, H., Torralba, A., Torresani, L. & Yan, Z. Hacs: Human action clips and segments dataset for recognition and temporal localization, 8668–8678 (2019).

[132] Gong, Z. et al. A large-scale fmri dataset for the visual processing of naturalistic scenes. Scientific Data 10, 559 (2023).

[133] Gifford, A. T., Cichy, R. M., Naselaris, T. & Kay, K. A 7t fmri dataset of synthetic images for out-of-distribution modeling of vision. arXiv preprint arXiv:2503.06286 (2025).

[134] Tucholka, A., Fritsch, V., Poline, J.-B. & Thirion, B. An empirical comparison of surface-based and volume-based group studies in neuroimaging. Neuroimage 63, 1443–1453 (2012).

[135] Marcus, D. S. et al. Human connectome project informatics: quality control, database services, and data visualization. Neuroimage 80, 202–219 (2013).

[136] Kay, K., Jamison, K. W., Zhang, R.-Y. & Uğurbil, K. A temporal decomposition method for identifying venous effects in task-based fmri. Nature methods 17, 1033–1039 (2020).

[137] Wu-Minn, H. 1200 subjects data release reference manual. URL https://www.humanconnectome.org 565, 2 (2017).

[138] Kay, K., et al. Disentangling signal and noise in neural responses through generative modeling. *bioRxiv* (2024).

[139] Wilcoxon, F. in Individual comparisons by ranking methods 196–202 (Springer, 1992).

